# Structural basis of substrate progression through the chaperonin cycle

**DOI:** 10.1101/2023.05.29.542693

**Authors:** Scott Gardner, Michele C. Darrow, Natasha Lukyanova, Konstantinos Thalassinos, Helen R. Saibil

## Abstract

The bacterial chaperonin GroEL-GroES promotes protein folding through ATP-regulated cycles of substrate protein binding, encapsulation, and release. Here, we have used cryoEM to determine structures of GroEL, GroEL-ADP·BeF_3_, and GroEL-ADP·AlF_3_-GroES all complexed with the model substrate Rubisco. Our structures provide a series of snapshots that show how the conformation and interactions of non-native Rubisco change as it proceeds through the GroEL-GroES reaction cycle. We observe specific charged and hydrophobic GroEL residues forming strong initial contacts with non-native Rubisco. Binding of ATP or ADP·BeF_3_ to GroEL-Rubisco results in the formation of an intermediate GroEL complex displaying striking asymmetry in the ATP/ADP·BeF_3_-bound ring. In this ring, four GroEL subunits bind Rubisco and the other three are in the GroES-accepting conformation, explaining how GroEL can recruit GroES without releasing bound substrate. Our cryoEM structures of stalled GroEL-ADP·AlF_3_-Rubisco-GroES complexes show Rubisco folding intermediates interacting with GroEL-GroES via different sets of residues.

## INTRODUCTION

Chaperonins prevent protein aggregation and promote correct folding through ATP-driven cycles of binding, encapsulation, and release of substrate proteins (1). The *E. coli* GroEL-GroES system is the archetypical chaperonin and is among the best studied molecular chaperones (2). GroEL subunits assemble into a tetradecameric complex composed of two back-to-back rings that surround a central cavity (3). The cavity is divided into two halves by the disordered C-terminal tails of GroEL subunits. Each GroEL monomer is divided into three domains: the nucleotide-binding equatorial domain, the apical domain that binds GroES and substrate, and an intermediate hinge domain. GroEL binds to non-native proteins through two apical domain helices (helices H and I) that form a hydrophobic collar around the entrance of each GroEL cavity (4). Binding of ATP causes conformational changes in GroEL that facilitate binding of the co-chaperonin GroES to seal the folding chamber (5). The ATP-induced conformational changes of GroEL can also lead to forced unfolding of the bound substrate, which occurs as a result of a stretching force applied to it (6). Forced unfolding of substrate proteins may be necessary to rescue them from kinetically trapped misfolded states and can enhance overall folding rates (7). However, a structural basis for forced unfolding by GroEL has not been described, and our previous cryoEM study of GroEL-ATP was carried out in the absence of a substrate protein (8).

Structural studies of different substrates bound to nucleotide-free GroEL have been published, including malate dehydrogenase (9), gp23 (10), PepQ (11), Rubisco (12), actin (13), and PrP (14). Together, these studies show that non-native proteins initially bind multivalently and in different configurations to GroEL apical domains, allowing GroEL to capture structurally distinct folding intermediates. The GroEL C-terminal tails, while not essential *in vivo*, also participate in capture and folding of substrate proteins, partly by promoting their deeper initial binding inside the GroEL cavity (15, 16).

Following binding of ATP and the heptameric co-chaperonin GroES, the GroEL-GroES cavity approximately doubles in volume and in principle can accommodate substrate proteins up to a mass of around 60 - 70 kDa (17). The majority of GroEL substrates identified in *E. coli* are 20 – 40 kDa, with a sharp cut-off toward proteins above 50 kDa (18). Two published cryoEM reconstructions of *Rhodospirillum rubrum* Rubisco (50.5 kDa) encapsulated by GroEL-GroES show either a native-like density (19) or a non-native-like density (15) located in the lower part of the cavity, interacting with hydrophobic residues of the cavity wall.

To gain insights into GroEL-assisted protein folding, we used cryoEM to determine structures of GroEL, GroEL-ADP·BeF_3_, and GroEL-ADP·AlF_3_-GroES, each complexed with the obligate substrate *Rhodospirillum rubrum* Rubisco. Our reconstruction of nucleotide-free GroEL-Rubisco shows that Rubisco fills the GroEL cavity and interacts with several GroEL apical domains and C-termini tails. By studying GroEL-ADP·BeF_3_-Rubisco, we identified a novel asymmetric conformation of the substrate-bound GroEL ring. Four GroEL subunits maintain contact with non-native Rubisco while the remaining three subunits extend upward. In addition, we observed a more extensive interaction between the substrate and the GroEL C-termini. This complex offers a possible mechanism for forced unfolding in which the substrate protein is stretched between the apical domains and C-termini of GroEL subunits while other GroEL subunits simultaneously present sites for GroES binding. Upon recruitment of GroES, Rubisco is encapsulated in the folding chamber where it adopts a native-like conformation held in place by interactions with hydrophobic and charged residues of GroEL-GroES.

## RESULTS

### CryoEM structure of GroEL bound to non-native Rubisco

Protein folding intermediates can be captured by rapidly diluting chemically denatured substrate proteins into a GroEL-containing buffer (20). We formed binary complexes of wild-type *E. coli* GroEL bound to the model substrate protein *R. Rubrum* Rubisco. We confirmed the formation of binary complexes using native gel electrophoresis and native mass spectrometry (Fig. S1).

Our initial attempts to determine a cryoEM reconstruction of GroEL-Rubisco were hindered by preferred orientation (Fig. S2). In the absence of non-native substrate, GroEL particles adopted a range of orientations permitting high resolution refinement (Fig. S2a). However, particles of GroEL bound to non-native Rubisco exhibited a strongly preferred end-view orientation. This limited the resolution of the reconstruction, and non-native Rubisco was not well resolved (Fig. S2b). To offset the preferred orientation, we collected cryoEM data employing stage tilt (Fig. S2c). Despite the lower number of particles, density for non-native Rubisco was better resolved (Fig. S2c).

To attain a higher resolution reconstruction, we aimed to reduce the interaction of GroEL-Rubisco with the air-water interface during cryoEM grid preparation. To accomplish this we prepared cryoEM grids of GroEL-Rubisco using a Chameleon (21, 22). Chameleon dispenses liquid sample onto a self-wicking grid, then after a short wicking time, plunge freezes the grid into liquid ethane. We collected cryoEM data of GroEL-Rubisco from two grids and determined a reconstruction from each. Although there was still preferred orientation, enough alternate views were recorded to yield isotropic reconstructions (Fig. S3a). We used 3D classification to remove apo GroEL particles then combined the GroEL-Rubisco particles from each data set (Fig. S4).

A 4.4 Å cryoEM map of GroEL-Rubisco was reconstructed from 65,453 particles (Fig. 1a, Fig. S3a, Fig. S4, and Table S1). The local resolution of the map ranged from 4.2 Å for GroEL equatorial domains to worse than 12 Å for non-native Rubisco (Fig. S3a). We refined the atomic model of GroEL (PDB code: 1SS8) into the cryoEM map (Fig. 1b). Extra density inside the GroEL cavities was attributed to bound non-native Rubisco. At a moderate contour level (5.0 σ), Rubisco density in the top ring represented the full volume estimated for a folded Rubisco monomer (∼61,000 Å^3^) (Fig. 1c). Density for non-native

Rubisco was also present in the bottom GroEL ring (Fig. 1a-c). However, this density was weaker and represented only around 30% of the volume of a folded Rubisco monomer (Fig. 1c). For comparison, the Rubisco density in reconstructions from our initial cryoEM data sets accounted for 20% - 50% of a natively folded monomer (Fig. S2).

Non-native Rubisco was positioned at the level of the GroEL apical domains (Fig. 1b). When displayed at a high contour level (8.0 σ), the map revealed individual contacts between non-native Rubisco and the apical domains of three GroEL subunits. Several additional features of the complex were apparent at lower map contour levels (< 5.0 σ) (Fig. 1c). At these lower contour levels, Rubisco density filled the top GroEL cavity and contacted all seven GroEL subunits via helix H, helix I, and the underlying hydrophobic segment. Weak interactions were observed between Rubisco and the C-terminal tails of several GroEL subunits (Fig. 1c). The Rubisco density also protruded ∼15 Å above the level of helix H. (Fig. 1c).

### Interactions between GroEL and non-native Rubisco

By thresholding the map through a range of contour levels (5 - 12 σ), we identified specific GroEL residues involved in binding non-native Rubisco (Fig. 1d). The strongest contacts to Rubisco were in helix I, consistent with previous cryoEM studies of different GroEL-substrate complexes (9–14). Many of the residues involved in contacting non-native Rubisco were those identified in the original mutational studies of GroEL (4). However, we also identified substrate-binding residues located at the C-terminal end of helix I, including M267, R268, and I270. The relative importance of these residues is less clear. GroEL-substrate interactions are canonically hydrophobic, explaining the contacts observed for M267 and I270. M267 has been implicated in allosteric communication, though an exact role is not known (23). Perhaps most surprising is the interaction with the positively charged residue R268, which was consistently the strongest interacting residue in our cryoEM reconstructions. GroEL R268 has previously been shown to form hydrogen bonds to glycine and serine residues on a 12-residue GroEL-binding peptide (24).

**Figure 1.**
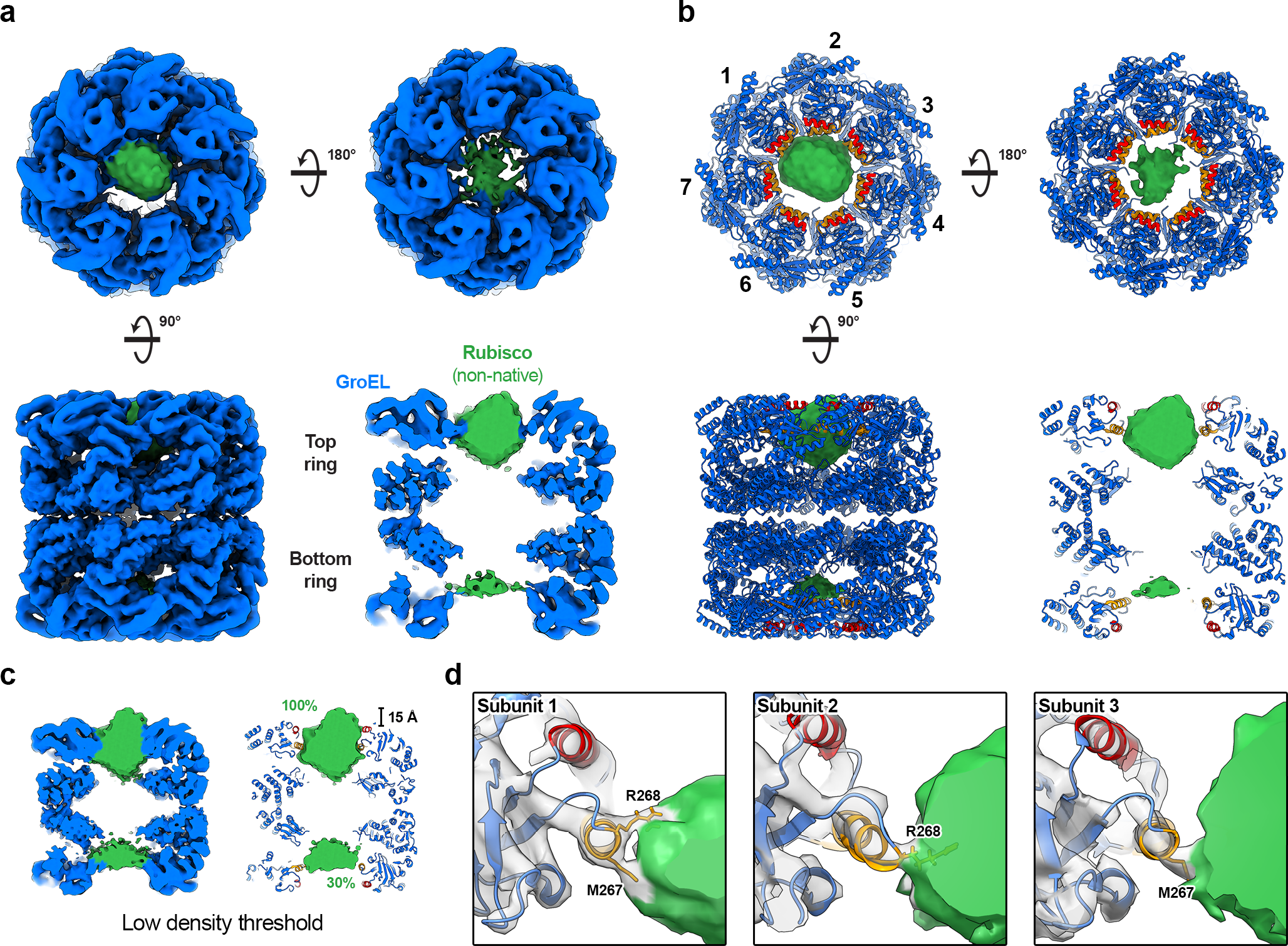
CryoEM structure of GroEL-Rubisco. **(a)** CryoEM map of GroEL-Rubisco at 4.5 Å. CryoEM density is shown coloured blue (GroEL) and green (Rubisco). **(b)** Refined atomic model of GroEL and Rubisco density (green) contoured at 8.0 σ. The atomic model of GroEL is coloured blue, the substrate-binding helices H and I are coloured red and orange respectively. **(c)** CryoEM map of GroEL-Rubisco contoured at a low threshold (5.0 σ). Percent values in green text represent the Rubisco density compared to that of a folded Rubisco monomer. **(d)** Contacts between GroEL subunits 1, 2, and 3 (grey density), and non-native Rubisco (green density). Interacting GroEL residues are labelled and shown as stick models.

### ATP binding induces asymmetry in the Rubisco-bound ring of GroEL

We next aimed to study the effects of ATP binding to GroEL-Rubisco, building on our previous work on GroEL-ATP (8). We formed GroEL-Rubisco complexes as described above, then added ATP several seconds before blotting and plunge-freezing using a Vitrobot. We collected cryoEM data employing stage tilt (25) to compensate for the preferred orientation of GroEL-ATP-Rubisco (Fig. S3b). Initial reconstructions showed that GroEL had partially denatured at the air-water interface (Fig. S5). We used a combination of signal subtraction and 3D classification to identify a subset of 13,015 relatively undamaged particles (Fig. S5). We determined a reconstruction of GroEL-ATP at a resolution of 4.3 Å (Fig. S5). The map showed a novel asymmetric ring arrangement of GroEL. However, the low resolution limited interpretability. We could not reliably identify bound nucleotide, and density for non-native Rubisco was not well resolved.

For high resolution cryoEM, we replaced ATP with a non-hydrolysable analogue. The ADP-metal complexes ADP·BeF_3_, ADP·AlF_3_, and ADP.VO_4_ are mimics of the ATP ground state, transition state, and post-hydrolysis state respectively (26). Both ADP·BeF_3_ and ADP·AlF_3_ support folding of the GroEL substrate Rhodanese in the presence of GroES (26). Both ATP analogues have been used to aid previous structural studies of chaperonins (10, 19, 27). Additionally, to reduce both preferred orientation and denaturation at the air-water interface, we used the Chameleon instrument to prepare grids for cryoEM.

A 3.4 Å cryoEM map of GroEL-ADP·BeF_3_-Rubisco was reconstructed from 202,582 particles (Fig. 2, Fig. S3c, Fig. S6, and Table S1). The map displayed an asymmetric ring and a symmetric ring (Fig. 2a). We observed density for Rubisco in the asymmetric ring only, contacting four GroEL subunits (Fig. 2a). The apical domains of the remaining three GroEL subunits were less well resolved and they extended upward, adopting a conformation reminiscent of the GroES-bound state (Fig. 2a). We used DeepEMhancer (28) to visualise the extended GroEL apical domains. However, Rubisco density was absent from the DeepEMhancer map, likely due to low local resolution. We used the locally filtered map from Relion to build a model of the complex and used the DeepEMhancer map only to position the apical domains of GroEL subunits 2, 5, and 7. We built and refined the model into the cryoEM maps using the crystal structures of apo GroEL (PDB code: 1SS8) and GroEL-GroES (PDB: 1SVT) (Fig. 2b). The conformation of the four substrate-contacting GroEL subunits resembled the Rs1 conformation previously reported for GroEL-ATP (Fig. 2c). In the Rs1 state, the GroEL intermediate and apical domains have undergone a 35° sideways tilt as a single rigid body relative to the nucleotide-free state (8). The four Rs1 GroEL subunits shared the equatorial-to-apical domain salt bridge, R58-E209, not observed in our previous study of GroEL-ATP (Fig. S3d). R58 is located within a short α-helix adjacent to the stem loop of GroEL equatorial domains. E209 lies in a short loop region of the underlying hydrophobic segment in the apical domains. In both the 2.7 Å crystal structure of apo GroEL (PDB: 1SS8) and in our 4.4 Å cryoEM reconstruction of nucleotide-free GroEL-Rubisco, the E209 loop faces away from the R58 helix and the R58-E209 sidechains are ∼8 Å apart (Fig. S3d). This salt bridge likely only forms upon binding of ATP (or analogue) and may act to stabilise the substrate-bound Rs1 state. The salt bridge could also be involved in allosteric communication between the apical and equatorial domains of GroEL. The residue E209 is located adjacent to the underlying hydrophobic segment which is involved in substrate-binding primarily via Y203. GroEL subunits 2, 5, and 7 adopted the GroES-bound state (Fig. 2d). This GroEL subunit conformation has only previously been observed in structures of GroEL-GroES, never in the absence of GroES. Our structure of GroEL-ADP·BeF_3_-Rubisco likely represents a transient intermediate complex adopted in response to ATP and substrate binding. This conformation of GroEL is presumably able to recruit GroES without releasing non-native substrate, and represents a missing link in substrate encapsulation.

**Figure 2.**
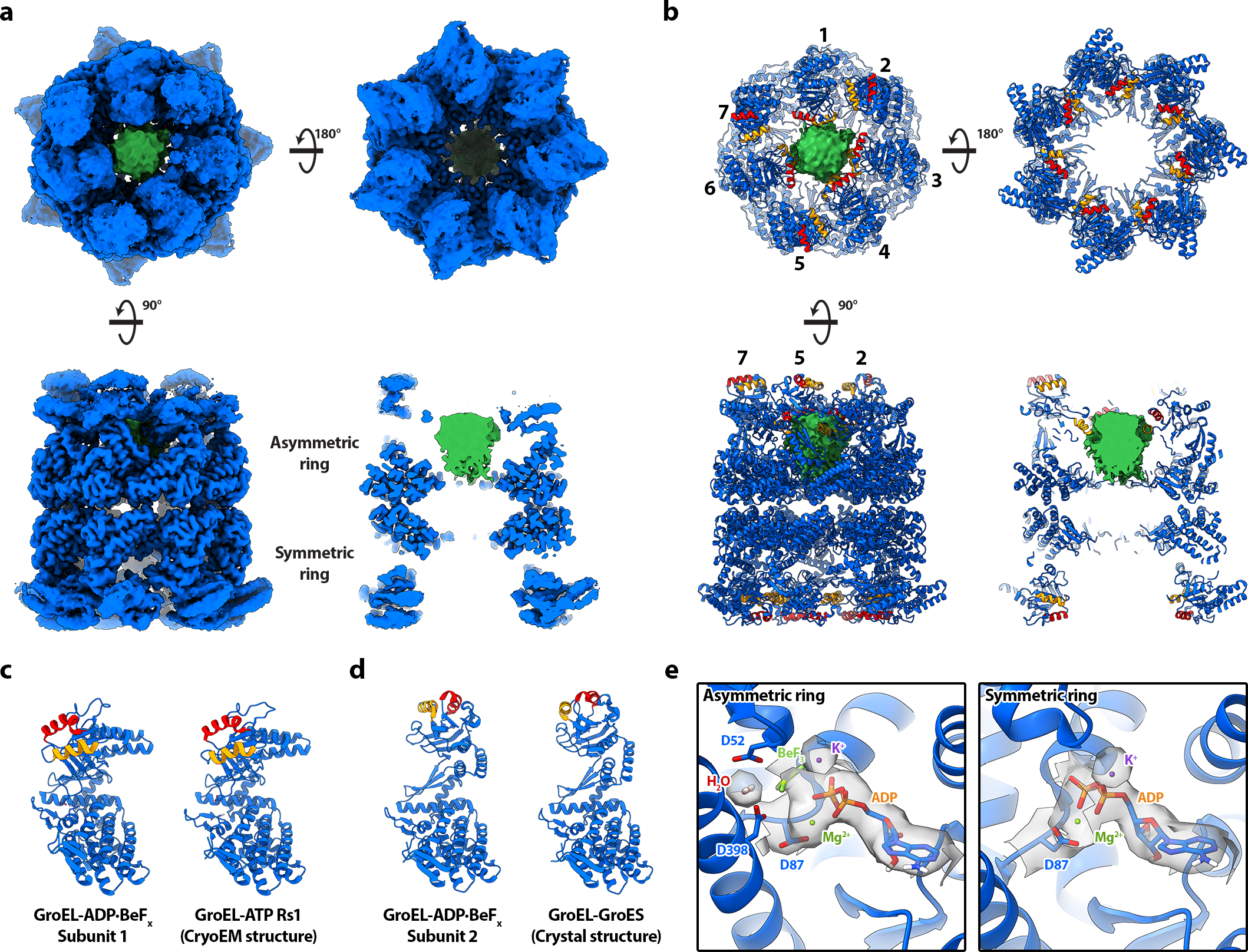
CryoEM structure of GroEL-ADP·BeF_3_-Rubisco. **(a)** CryoEM map of GroEL-ADP·BeF_3_-Rubisco at 3.4 Å. The GroEL map (blue) displayed was generated by DeepEMhancer. Density for non-native Rubisco (green) was isolated from the locally filtered map generated by Relion. **(b)** Refined atomic model of GroEL-ADP·BeF_3_ and non-native Rubisco density (green). The substrate-binding helices H and I are coloured red and orange respectively. The asymmetry can be appreciated from the position of helix H in each subunit. **(c)** Comparison of GroEL-ADP·BeF_3_ subunit 1 with the published structure of the Rs1 conformation of GroEL-ATP (PDB: 4AAQ). **(d)** Comparison of GroEL-ADP·BeF_3_ subunit 2 with the published crystal structure of GroEL-GroES (PDB: 1SVT). **(e)** Nucleotide binding sites of each GroEL ring, showing ADP·BeF_3_ in asymmetric ring subunits and ADP in symmetric ring subunits. Overlaid cryoEM density is shown only for the labelled moieties.

We examined the nucleotide binding sites of GroEL subunits (Fig. 2e). Clear density for ADP was seen in all fourteen sites. We observed differences in nucleotide site density between the two rings, but saw no obvious differences among subunits belonging to the same ring. We observed continuous density between the GroEL D87 sidechain and ADP. D87 is involved in ATP hydrolysis, and mutations such as D87K abolish ATPase activity (4). We were able to confidently model ADP and the phosphate oxygen-coordinating metal, Mg^2+^. ADP bound in the asymmetric ring showed additional density that we attributed to the ATP γ-phosphate analogue, BeF_3_. (Fig. 2e). Symmetric ring ADP lacked this additional density and it was modelled without BeF_3_. Subunits in both rings showed density for the second coordinating metal ion, K^+^. GroEL requires K^+^ to hydrolyse ATP, and a previously published crystal structure confirmed this position as the K^+^ binding site (29). In the asymmetric ring, we observed additional density between the D52 and D398 sidechains (Fig. 2e). We attributed this to the water molecule involved in attacking the γ-phosphate of ATP during hydrolysis (30). Asymmetric ring subunits have therefore been captured in an ATP-bound state prior to hydrolysis, consistent with the classification of ADP·BeF_3_ as a ground-state analogue of ATP.

### Interactions between non-native RuBisCO and GroEL-ADP·BeF3

Non-native Rubisco interacted with the apical domains of four GroEL subunits in the asymmetric ring (Fig. 3). At low contour levels, the interaction was dominated by helix I and the underlying hydrophobic segment of GroEL subunits 1, 3, 4 and 6 (Fig. 3a), leaving subunits 2, 5, and 7 to extend upward. Rubisco was located deeper in the GroEL cavity than in our cryoEM reconstruction of nucleotide-free GroEL-Rubisco, and did not protrude above the level of helix H. At a moderate contour level (5.0 σ), the volume of Rubisco density in the asymmetric ring represented ∼68% of a full Rubisco monomer. The map also showed non-native Rubisco forming contacts to the C-termini of all seven GroEL subunits (Fig. 3a). In the symmetric ring, we observed density for the C-terminal GroEL residues P525 and K526 (typically disordered in crystal structures), but no density for non-native Rubisco (Fig. 3a). We examined the contacts between GroEL apical domains and non-native Rubisco at higher contour levels (Fig. 3b). The strongest interactions were similar to those we observed in the nucleotide-free binary complex (Fig. 1). The same R268 contact in helix I was seen for each of the four GroEL subunits (Fig. 3b). Other subunits showed contacts involving V264 and N265 of helix I, and residue Y203 of the underlying hydrophobic segment (Fig. 3b).

**Figure 3.**
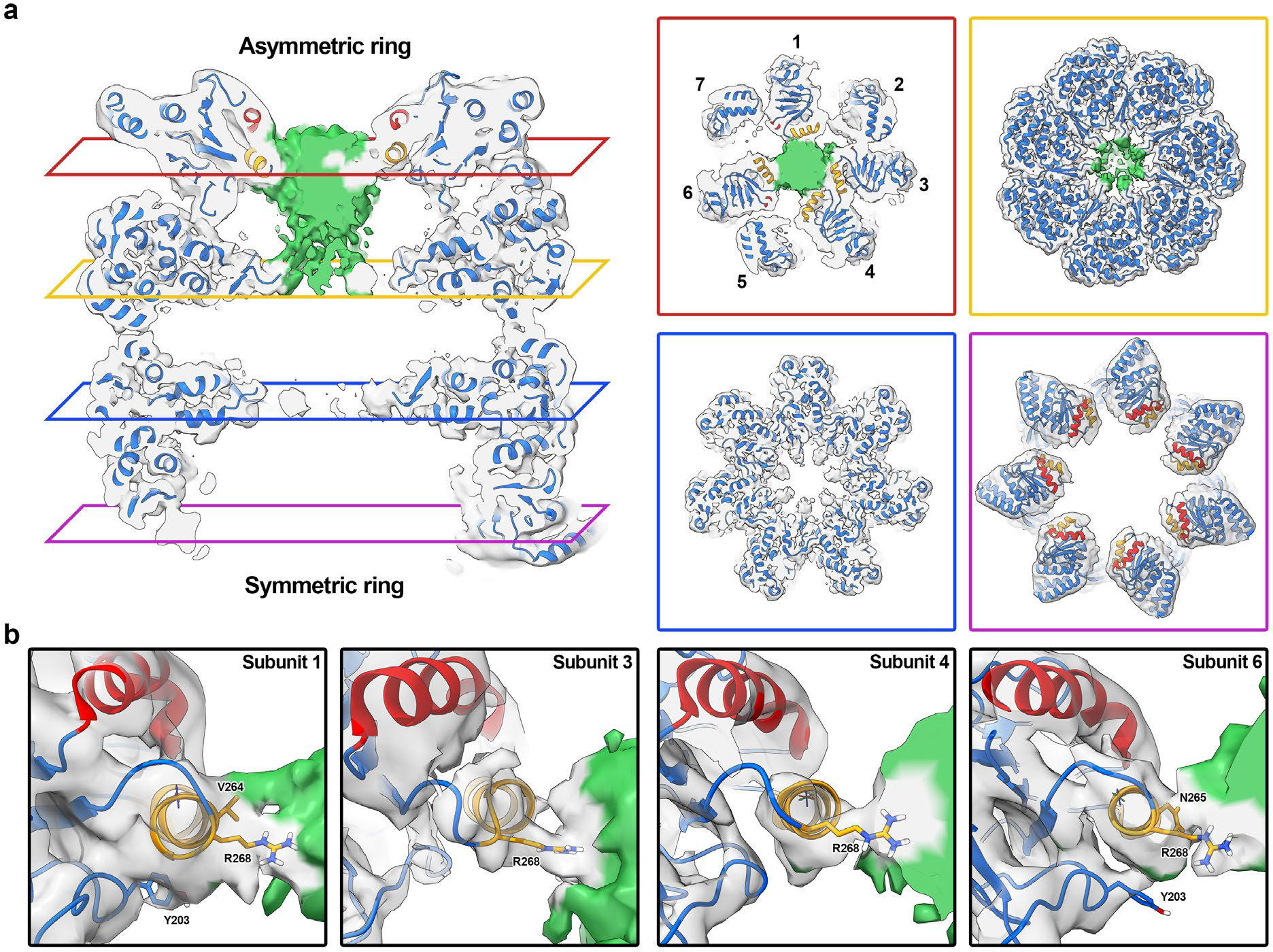
Interactions between GroEL-ADP·BeF_3_ and non-native Rubisco. **(a)** Central slices through the GroEL-ADP·BeF_3_-Rubisco model overlaid with the cryoEM map (transparent grey). Panels showing lateral slices through the asymmetric ring apical domains (red panel), asymmetric ring equatorial domains (yellow panel), symmetric ring equatorial domains (blue panel), and symmetric ring apical domains (purple panel). **(b)** Interactions between GroEL apical domains and non-native Rubisco.

### Rubisco encapsulated by GroEL-ADP·AlF3-GroES

We next studied the conformation of encapsulated Rubisco in the full GroEL-GroES complex. We added GroES and ADP·AlF_3_ to GroEL-Rubisco to form stalled ternary complexes. We again used a Chameleon instrument to prepare frozen grids. Initial 3D classification showed variability in the occupancy of GroES (Fig. S7). Two of the 3D classes showed GroEL-ADP·AlF_3_ complexes without GroES. Since reconstructions from these classes resembled our structure of GroEL-ADP·BeF_3_, we discarded these particles. To identify particles with encapsulated Rubisco, we used masked 3D classification targeting the *cis* cavity of GroEL-GroES (Fig. S7). We determined a 3.7 Å reconstruction of GroEL-ADP·AlF_3_-Rubisco-GroES from 30,965 particles (Fig. 4a, Fig. S3e, Fig. S7, and Table S1). We refined the published crystal structure of GroEL-GroES (PDB: 1SVT) into our cryoEM map (Fig. 4b). Density for encapsulated Rubisco occupied the upper two-thirds of the *cis* cavity, adjacent to the GroEL apical domains. The Rubisco density accounted for 40 - 50 kDa of protein mass, and its shape was reminiscent of a folded Rubisco monomer. Interactions were observed with several cavity-facing residues of GroEL-GroES subunits (Fig. 4c). The strongest contacts to encapsulated Rubisco involved GroEL residue F281, and GroES residue Y71 (Fig. 4c). The Y71 residue of GroES subunits form a hydrophobic collar on the ceiling of the *cis* cavity and may be important for the folding of some GroEL substrates (31). Additional contacts were resolved at lower map contour levels and involved GroEL residues R197, K226, N229 and E255 (Fig. 4c, right panel).

**Figure 4.**
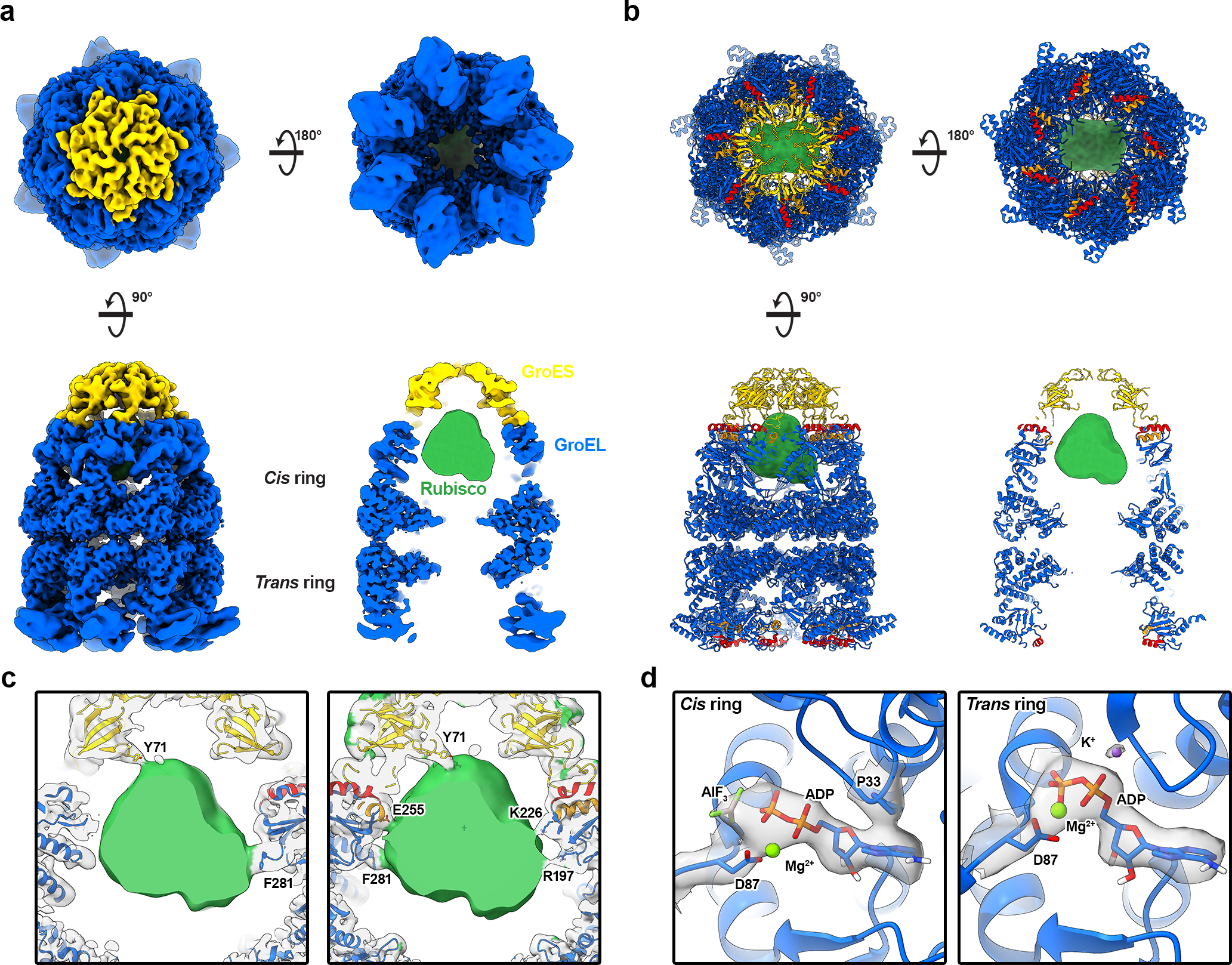
CryoEM structure of GroEL-ADP·AlF_3_-Rubisco-GroES. **(a)** CryoEM map of GroEL-ADP·AlF_3_-Rubisco-GroES at a global resolution of 3.7 Å, filtered by local resolution. **(b)** Refined atomic model of GroEL-ADP·AlF_3_-GroES and non-native Rubisco density (green). **(c)** Molecular contacts between GroEL-GroES and Rubisco. **(d)** Nucleotide site density in the *cis* and *trans* rings.

The conformation of the *trans* ring of GroEL-ADP·AlF_3_-Rubisco-GroES resembled the symmetric ring of GroEL-ADP·BeF_3_-Rubisco. This “wide” conformation of the GroEL *trans* ring is likely related to the presence of high concentrations of ADP (3 mM) during sample preparation (32). We did not observe density for non-native Rubisco in the *trans* ring, even at low map contour levels. In this conformation, the continuous hydrophobic collar formed by helices H and I is disrupted, possibly leading to reduced substrate binding. Lower concentrations of ADP may have allowed for visualisation of bound non-native substrate in the *trans* ring.

We observed clear density for ADP in all fourteen nucleotide binding sites. *Cis* ring ADP showed additional density that we modelled as AlF_3_ (Fig. 4d). *Trans* ring sites contained the coordinating potassium ion (Fig. 4d). At this point along the ATP hydrolysis reaction coordinate, K^+^ has presumably fulfilled its catalytic role and is no longer required in the *cis* ring. In contrast, our structure of GroEL-ADP·BeF_3_-Rubisco showed K^+^ bound in the ADP·BeF_3_-bound ring, but not in the ADP-bound ring. This is consistent with BeF_3_ and AlF_3_ mimicking different states of the ATP γ-phosphate.

### Further 3D classification revealed distinct conformations of encapsulated Rubisco

The Rubisco density in our reconstruction of the stalled ternary GroEL-ADP·AlF_3_-Rubisco-GroES complex likely represented an ensemble of conformations that had been averaged together during image processing. We aimed to identify some of these conformations using an additional round of 3D classification (Fig. S7). Due to the relatively low number of particles at this processing step, we opted to use four classes for masked 3D classification, targeting the GroEL-GroES *cis* cavity. We refined each subset of particles to a resolution of 4.1 - 4.2 Å (Fig. S3f) and performed a rigid body fit of our refined model of GroEL-ADP·AlF_3_-GroES (Fig. 5). In the four reconstructions the interactions between GroEL-GroES and the encapsulated Rubisco monomer were well resolved. Each class showed a different set of GroEL-GroES residues interacting with Rubisco, suggesting that GroEL-GroES can stabilise a range of non-native substrate conformations (Fig. 5). At lower contour levels the substrate density accounted for the volume a full Rubisco monomer (∼61,000 Å^3^). All four classes shared the same GroEL F281 contact to Rubisco (Fig. 5a-d, red dashed circles). Individual classes displayed additional contacts from the Rubisco density to GroEL residues K226 (Fig. 5a), N229 (Fig. 5a), E255 (Fig. 5c-d), and Y360 (Fig. 5c). Additionally, class 4 displayed strong contacts to the Y71 residues of two adjacent GroES subunits (Fig. 5d).

**Figure 5.**
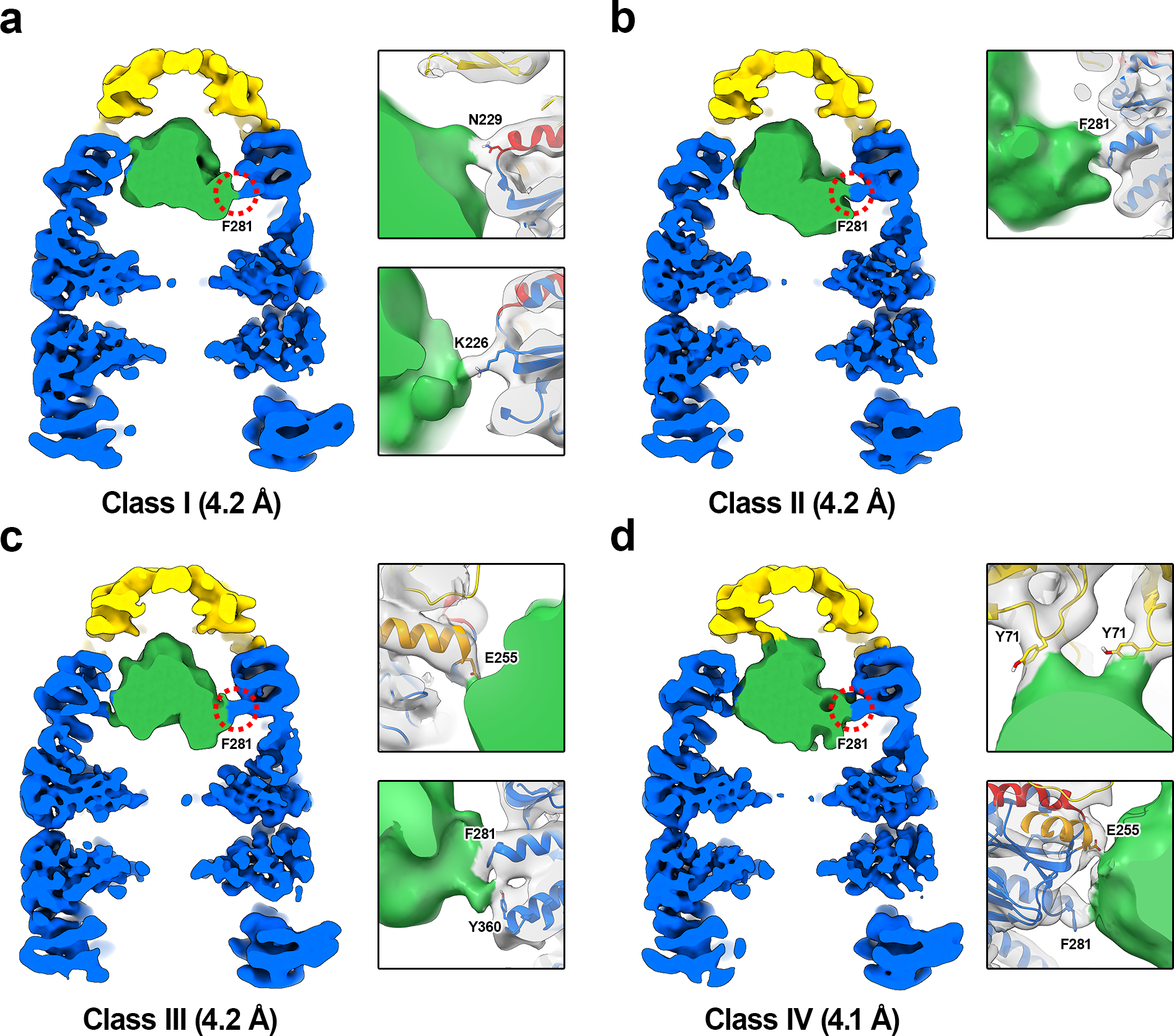
Multiple classes of encapsulated Rubisco. Red dashed circles highlight the contact in all four reconstructions between GroEL residue F281 and Rubisco. **(a)** Reconstruction of class I from 7,202 particles. Panels highlight the K226 and N229 contacts. **(b)** Reconstruction of class II from 8,237 particles. Panel highlights the F281 contact. **(c)** Reconstruction of class III from 7,818 particles. Panels highlight the F281, Y360, and E255 contacts. **(d)** Reconstruction of class IV from 7,708 particles. Panels highlight the E255, F281 and GroES Y71 contacts.

### Model of near-native Rubisco encapsulated in the GroEL-GroES folding chamber

To model Rubisco we examined the density in the four cryoEM reconstructions. Due to the low local resolution, we were unable to identify secondary structure elements of Rubisco. We limited our analysis to low resolution features and examined density that might represent the different Rubisco domains. Rubisco monomers are comprised of two domains, an N-terminal domain (NTD; residues 1 – 135) and a larger C-terminal TIM barrel domain (CTD; residues 136 – 466) (33). Classes I, III, and IV could accommodate rigid body fits of the Rubisco monomer in multiple different orientations. The class II reconstruction (Fig. S7) showed two distinct lobes of density when displayed at a higher contour level (Fig. 6a). We attributed these lobes to the NTD and CTD of Rubisco and used them to orient and rigid body fit the published Rubisco crystal structure (PDB: 9RUB) (Fig. 6b). We flexibly fit the Rubisco monomer into the density, allowing for only minor changes when optimising the map-model fit (Fig. 6b-c and Table S1).

**Figure 6.**
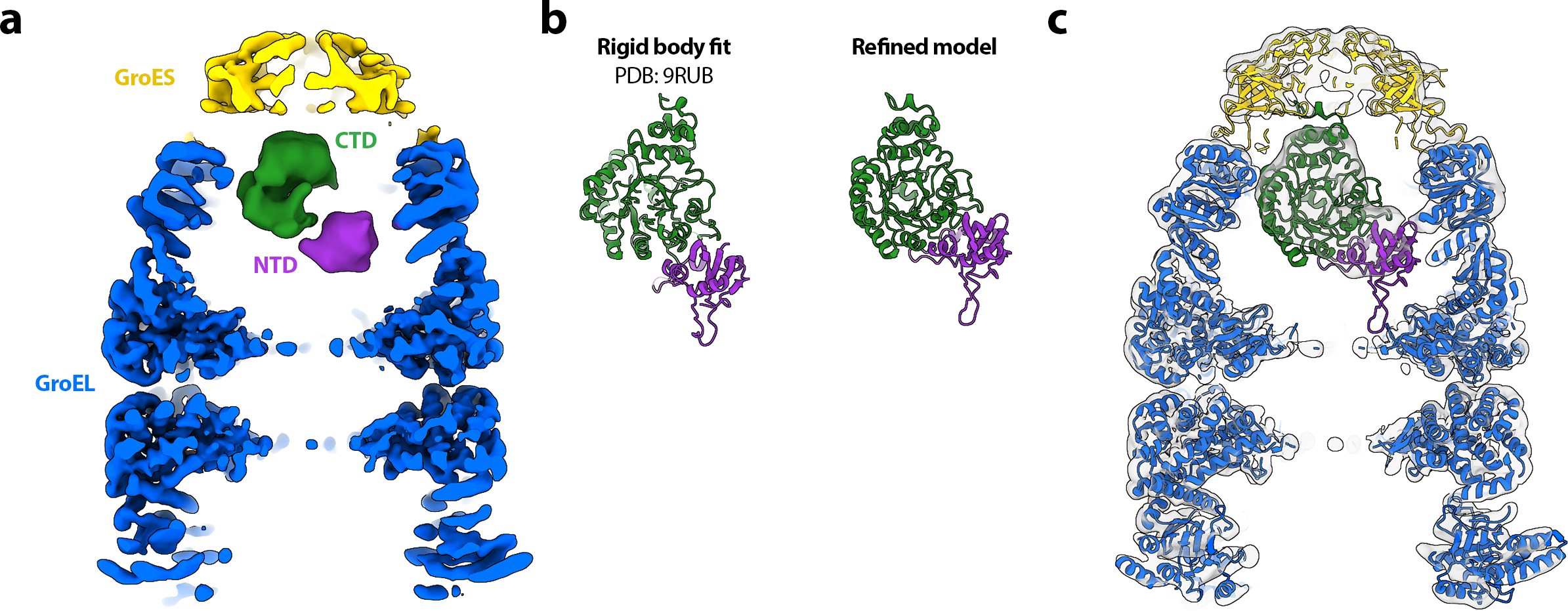
Modelling Rubisco inside the GroEL-GroES folding chamber. **(a)** CryoEM map of GroEL-ADP·AlF_3_-Rubisco-GroES (class II) at a contour level of 7σ. The two domains of the encapsulated Rubisco monomer are coloured purple (NTD) and green (CTD). **(b)** Comparison between the crystal structure of a Rubisco monomer and the refined model. **(c)** Refined model of GroEL-ADP·AlF_3_-Rubisco-GroES overlaid on the class II density at a contour level of 3σ.

## DISCUSSION

In this study, we have used single-particle cryoEM to determine structures GroEL, GroEL-ADP·BeF_3_ and GroEL-ADP-AlF_3_-GroES all complexed with non-native Rubisco. Our work provides a series of snapshots of a non-native protein as it progresses through the GroEL-GroES reaction cycle, revealing the interactions between GroEL and substrate at each step. We have described a novel conformation of ATP-bound GroEL that can simultaneously recruit its co-chaperonin GroES while still binding non-native substrate, preventing its escape during the encapsulation step. Lastly, we showed that encapsulated Rubisco resides in the GroEL-GroES cavity as an ensemble of conformational states that likely represent different folding intermediates.

We have previously shown that Rubisco binds to the apical domains of GroEL subunits (12) in a similar fashion to other GroEL substrates (9–11, 13, 14). Structural studies of GroEL-substrate complexes are typically limited to low resolution and density for non-native substrate is usually incomplete. Our cryoEM reconstruction of GroEL-Rubisco showed that the strongest contacts were formed with helix I of GroEL, consistent with previous structural studies of other GroEL-substrate complexes. Several of the residues involved in contacting non-native Rubisco were those identified in the original mutational studies of GroEL, such as V264 and Y203 (4). Recognition of substrates by GroEL is typically described as predominantly hydrophobic. However, the strongest contact to Rubisco involved the GroEL residue R268, located at the C-terminus of helix I. The importance of R268 in substrate binding and folding is less well characterised. Structural evidence for the role of R268 in mediating substrate interactions comes from the crystal structure of GroEL bound to a 12-residue peptide (24). The GroEL-peptide structure showed that a serine and glycine residue on the peptide formed hydrogen bonds to R268 (24). Our cryoEM structures of GroEL-Rubisco and GroEL-ADP·BeF_3_-Rubisco identified R268 as an important residue in mediating non-native substrate interactions prior to GroES binding. Additionally, we observed a strong interaction between non-native Rubisco and GroEL residue M267, located on the lower face of helix I. To our knowledge, M267 has not previously been implicated in substrate binding, though it may play a role in intra- and inter-subunit allosteric communication (23).

Experiments using native mass spectrometry have previously shown that Rubisco monomers bind to GroEL tetradecamers with a 1:1 stoichiometry, exerting negative cooperativity on the opposite GroEL ring and inhibiting binding of a second Rubisco monomer (34). Other substrates with a range of molecular weights (32 – 56 kDa) have been shown to bind to both GroEL rings simultaneously, implying that GroEL can recognise and respond to different types of substrate. (35). Our native mass spectrometry results for GroEL-Rubisco agreed with the published results. However, our cryoEM reconstruction showed Rubisco bound in both rings simultaneously, albeit with different occupancies. Previous work has suggested that the structural basis for this negative cooperativity lies in a narrowing of the opposite GroEL ring (12, 15, 35). However, we did not observe a significant structural change in the opposite ring.

All previously published cryoEM structures of GroEL-substrate complexes were determined from grids prepared using conventional plunge-freezing methods that included a blotting step, and a several-second delay between sample application and vitrification. It has been shown that reducing this delay can reduce denaturation at the air-water interface and improve the orientation distribution of particles (22). Previous reconstructions of GroEL-substrate complexes did not display the full expected volume of the non-native substrate. For example, previous studies report the following percentages for the substrate volume in their reconstruction; GroEL-MDH: 25 - 40% (9), GroEL-gp23: 54% (10), GroEL-actin: 28% (13), and GroEL-Rubisco: 30 - 35% (12). Our initial cryoEM attempts using traditional vitrification methods yielded similar results, but our reconstruction from Chameleon grids accounted for the full volume of Rubisco. It is likely that non-native substrate in previous studies had been partially denatured at the air-water interface. The missing density in the published reconstructions would have presumably protruded from the GroEL cavity, as displayed in our reconstruction of GroEL-Rubisco. Non-native proteins are particularly prone to adsorption at the air-water interface during cryoEM grid preparation. Our work shows that reducing the time between sample application and vitrification provides additional benefits in the study of biological systems involving non-native proteins.

Binding of ATP is known to trigger conformational changes within GroEL subunits (8, 36). Studies of GroEL-GroES bound to ATP analogues such as ATPγS or AMP-PNP fail to form folding active complexes and do not show the same large-scale conformational changes in GroEL (26, 37, 38). This is likely related to the critical role of the ATP γ-phosphate, mimicked in our structures by BeF_3_ or AlF_3_ (26). Here, we present the first structure of a nucleotide-bound and substrate-bound GroEL complex.

Our previous structural study of GroEL-ATP was carried out in the absence of substrate protein, and the data set was not large enough to test for asymmetry, particularly in the more open states (8). In this work, we show that individual GroEL subunits of the ATP-bound ring adopt one of two conformations, resulting in a markedly asymmetric ring. This asymmetric behaviour of subunits is reminiscent of the eukaryotic hetero-oligomeric group II chaperonin TRiC/CCT (39). Our cryoEM structure of GroEL-ADP·BeF_3_-Rubisco reveals how some GroEL subunits can recruit GroES while others are bound to non-native substrate, preventing its escape. The four substrate-bound GroEL subunits adopt the Rs1 state reported for GroEL-ATP (8). The three remaining subunits adopt the GroES-bound conformation (40), despite the absence of GroES itself.

In our cryoEM map of GroEL-ADP·BeF_3_-Rubisco, the C-terminal tails of all seven asymmetric ring subunits contacted non-native Rubisco. In comparison, structures of nucleotide-free GroEL-substrate complexes do not typically suggest an extensive interaction with the C-termini. This suggests that deeper substrate-binding role of the GroEL C-termini becomes more important following ATP binding. Deletion of the GroEL C-termini has been shown to slow folding of Rubisco (16). Nevertheless, the C-termini themselves are not essential *in vivo* (41) despite being highly conserved among chaperonins. The slowed rate of folding upon C-termini tail deletion has been attributed to altered rates of chaperonin cycling and ATPase activity (discussed in appendix 1 of the comprehensive review of chaperonins in ref (2)).

We previously speculated that GroES is initially recruited by 1 - 2 raised GroEL subunits, resulting in an asymmetric intermediate (8, 9). However, significant asymmetry of a GroEL ring has only been previously reported in the crystal structure of the double mutant GroEL_ΔD83A/ΔR197A_ bound to ADP (42). In this mutant, two inter-subunit salt bridges were removed, effectively detaching adjacent apical domains. The freed apical domains adopted conformations similar to those observed for GroEL-ATP (8). Our structure of GroEL-ADP·BeF_3_-Rubisco offers the first view of an asymmetric wild-type GroEL ring.

During transition from the Rs1 state to the GroES-bound state, GroEL apical domains undergo a dramatic upward swing of 60°, and a 90° clockwise rotation (8). These movements have been suggested to exert a stretching force on the substrate, which remains bound to several apical domains during their motion (43). This stretching action is thought to forcefully unfold GroEL substrates, rescuing kinetically trapped folding intermediates (6, 16). Our previous study of GroEL-ATP suggested a possible mechanism for forced unfolding in which the radial expansion of subunits exposed bound substrate to stretching (8). Our structure of GroEL-ADP·BeF_3_-Rubisco suggests a different geometrical pathway of stretching. A substrate that is multivalently bound between GroEL apical domains and C-termini could be stretched during their transition from Rs1 to GroES-bound states. In support of this, it has previously been shown that forced unfolding of Rubisco is attenuated when the C-termini are removed (16). Bound Rubisco might also be destabilised due to the exclusion of bulk water from the occupied GroEL cavity, reducing the hydrophobic effect and altering the energetics of its folding relative to that in bulk solution (44). Following GroES binding, non-native Rubisco would be released into the folding chamber where it may still associate with the C-termini (19).

Our reconstructions of GroEL-ADP·AlF_3_-Rubisco-GroES showed a native-like Rubisco in the upper half of the folding chamber, held in place by interactions with charged and hydrophobic residues located in the GroEL apical domains (K226, E255, F281, Y360), and in GroES (Y71). Several cryoEM structures of GroEL-GroES-substrate complexes, including two with Rubisco as the substrate protein, have been published (10, 15, 19). Importantly, the published structure of GroEL_43Py398A_-GroES (15) is not a fully folding-active complex and instead represents a stalled complex immediately prior to the release of the substrate inside the folding chamber. Both previously published structures of GroEL-ADP·AlF_3_-Rubisco-GroES showed either non-native or native-like Rubisco located in the lower half of the folding chamber, interacting with residues of the GroEL apical domains (F281, Y360), the equatorial domains (F44), and the GroEL C-termini (15, 19). GroEL residue F281 appears to be critical in the folding of substrate proteins. This is supported by earlier work showing that the mutant GroEL_F281D_ supports binding of non-native substrate, but exhibits decreased ATPase activity, reduced folding, and aggregation of the substrate protein upon its release (4).

The position of Rubisco in GroEL-GroES was similar position to that of the T4 bacteriophage capsid protein, gp23 (10), also observed in a native-like state. Rubisco interacted with several of the GroES Y71 residues that together form a hydrophobic ring on the folding chamber ceiling. Previous work has shown that this hydrophobic ring may take part in the folding process for some GroEL substrates (31). We did not observe contacts with hydrophobic residues in the lower part of the chamber, such as F44 or the C-termini, and instead observed interactions with charged residues in the GroEL apical domains. Encapsulated substrates may start as folding intermediates at the bottom of the GroEL-GroES cavity, sequestered primarily by the C-termini and the F44 loop. As folding proceeds and hydrophobic residues in the substrate become buried, the interaction with the C-termini might diminish, allowing the substrate to occupy a more central or upper position in the cavity. The Rubisco intermediate in our structure likely represents a near-native state, with some distortion of the domain interface, primed for release following detachment of GroES.

## CONCLUSION

Our results, benefitting from the substantial advances in cryoEM methodology in the last decade, provide a new, more detailed view of the chaperonin assisted folding pathway and mechanism for a major, model substrate. Our cryoEM reconstructions show the changing sites of interaction on GroEL and then GroES during folding, and progression through key initial steps in the nucleotide cycle, as well as displacement in the GroEL-GroES cavity as native structure is formed.

## METHODS

### Protein expression and purification

#### Escherichia coli GroEL

Plasmid pTrcESL, a kind gift from Dr. Peter Lund (University of Birmingham, UK), was transformed into BL21 E. coli. Cells were grown in LB media at 37 °C using baffled flasks. Overexpression of GroELS was induced by addition of 0.5 mM isopropyl β-D-thiogalactopyranoside (IPTG) at an A600 of 0.6 - 0.8. After 3.5 hours cells were harvested by centrifugation, resuspended in chilled lysis buffer (50 mM Tris, pH 7.4), and disrupted using an EmulsiFlex-C3 (Avestin, Ottawa, Canada). Cellular debris and insoluble material were removed by centrifugation at 55,000 RCF at 4 °C for 1 hour. Subsequent purification and dialysis steps were performed at 4 °C unless stated. Soluble lysate was filtered through a 0.45 µm syringe filter and loaded onto a 140 mL Q Sepharose FastFlow ion-exchange column (GE Healthcare) equilibrated in buffer A (50 mM Tris, pH 7.4, 1 mM DTT). The column was washed with 20% buffer B (50 mM Tris, pH 7.4, 1 mM DTT, 1M NaCl) until A280 reached a steady baseline. Protein was eluted with a 20 - 50% 15 column volume gradient of buffer B and fractions were collected. Elution of GroEL was confirmed by SDS-PAGE and fractions corresponding to the latter half of the GroEL peak were pooled. Solid ammonium sulphate was added slowly to the pooled fractions to a final concentration of 1.2 M and stirred overnight. Pooled fractions from ion-exchange chromatography were filtered through a 5 µm syringe filter and loaded onto a 22 mL Source 15ISO hydrophobic interaction column (GE Healthcare) equilibrated in buffer C (50 mM Tris, pH 7.4, 1 mM DTT, 1.2 M NH4(SO4)2. The column was washed with buffer C until the A280 reached a steady baseline < 50 mAU. GroEL was eluted with a 14 column volume reverse-gradient of 100 - 0% buffer C to buffer A. Peak fractions were analysed by SDS-PAGE. The protein concentration and tryptophan fluorescence of each peak fraction were measured. Since GroEL has no tryptophan residues, tryptophan fluorescence is an indicator of contamination by bound Trp-containing proteins. Tryptophan fluorescence was measured using a FluoroMax 3 (Horiba). Protein concentration was measured by Pierce BSA protein assay. Fractions with high protein concentration and low fluorescence were pooled and dialysed overnight against 4 L of buffer D (50 mM Tris, pH 7.4, 50 mM KCl, 1 mM DTT). The final peak fractions from hydrophobic interaction chromatography typically yielded the purest GroEL. To further strip GroEL-bound contaminating proteins, the dialysed protein was concentrated to 20 mg/mL, then 2 mL of buffer E (1:1 mixture of buffer D and Affi-gel Blue resin) was added per 30 mg of total protein. Methanol was then added dropwise to 20% v/v while stirring. After mixing, the sample was rocked gently at room temperature (∼ 21 °C) for 2 hours. GroEL was recovered in the supernatant by filtration through a 1.2 µm syringe filter and the resin was washed once with its volume of buffer D. Eluates were pooled and dialysed overnight against 4 L of buffer D. Purified GroEL was filter-sterilised and concentrated to 30 - 40 mg/mL. Aliquots of concentrated GroEL were flash frozen in liquid nitrogen and stored at −80 °C until use.

#### Escherichia coli GroES

Plasmid pTrcESL was transformed into BL21 E. coli. Cells were grown in LB media at 37 °C using baffled flasks. Overexpression of GroELS was induced by addition of 0.5 mM IPTG at an A600 of 0.6 - 0.8. After 3.5 hours cells were harvested by centrifugation, resuspended in chilled lysis buffer (50 mM Tris, pH 7.4), and disrupted using an EmulsiFlex-C3 (Avestin, Ottawa, Canada). Cellular debris and insoluble material were removed by centrifugation at 55,000 RCF at 4 °C for 1 hour. Soluble lysate was transferred to a fresh tube and acidified to pH 5.0 by dropwise addition of 1 M acetic acid. Precipitated material was removed by centrifugation at 25,000 RCF at 4 °C for 15 min. Subsequent purification and dialysis steps were performed at 4 °C unless stated. Acidified soluble lysate was filtered through a 0.45 µm syringe filter and loaded onto a 125 mL SP Sepharose FastFlow ion-exchange column (GE Healthcare) equilibrated in buffer A (50 mM NaOAc, pH 5.0). The column was washed with buffer A until A280 reached a steady baseline. Protein was eluted with a 0 - 50% 12.5 column volume gradient of buffer B (50 mM NaOAc, pH 5.0, 1 M NaCl) and fractions were collected. Elution of GroES was confirmed by SDS-PAGE. Fractions that corresponded to the GroES peak were pooled, neutralised to pH 7.5 - 8.0 with Tris-base, and dialysed overnight against 4 L of buffer C (50 mM Tris, pH 7.4). Dialysed protein was loaded onto a 24 mL Source 15Q ion-exchange column equilibrated in buffer C. The column was washed with buffer C until the A280 reached a steady baseline < 50 mAU. GroES was eluted with a 0 - 50% 20 CV gradient of buffer D (50 mM Tris, pH 7.4, 1 M NaCl). Elution of GroES was confirmed by SDS-PAGE. Fractions that corresponded to GroES were pooled and dialysed overnight against 4 L buffer C. Purified GroES was filter-sterilised and concentrated to 15 mg/mL. Aliquots of concentrated GroES were flash frozen in liquid nitrogen and stored at −80 °C until use.

#### Rhodospirillum rubrum RuBisCO

Plasmid pT7Rub, a kind gift from Dr. Wayne Fenton (Yale University, USA), was transformed into BL21 (DE3) E. coli. Cells were grown in LB media in baffled flasks at 20°C for 24 hours in the absence of induction. Cells were harvested by centrifugation, resuspended in lysis buffer (50 mM Tris, pH 7.4), and disrupted using an EmulsiFlex-C3 (Avestin, Ottawa, Canada). Cellular debris and insoluble components were removed by centrifugation at 55,000 RCF at 4 °C for 1 hour. Subsequent purification and dialysis steps were performed at 4 °C unless stated. Soluble lysate was filtered through a 0.45 µm syringe filter and loaded onto a 140 mL Q Sepharose FastFlow ion-exchange column (GE Healthcare) equilibrated in buffer A (50 mM Tris, pH 7.4, 1 mM DTT). The column was washed with buffer A (50 mM Tris, pH 7.4, 1 mM DTT) until A280 reached a steady baseline. Rubisco was eluted with a 0 - 50% 10 column volume gradient of buffer B (50 mM Tris, pH 7.4, 1 mM DTT, 1M NaCl) and fractions were collected. Elution of Rubisco was confirmed by SDS-PAGE and peak fractions were pooled and dialysed overnight against 4L of buffer C (20 mM Tris, pH 7.4, 1 mM DTT). Pooled fractions from ion-exchange chromatography were filtered through a 1.2 µm syringe filter and loaded onto a 60 mL Affi-Gel Blue column equilibrated in buffer C. The column was washed with buffer C until the A280 reached a steady baseline. Rubisco was eluted with a 0 - 50% 4 column volume gradient of buffer D (20 mM Tris, pH 7.4, 1 mM DTT, 1M NaCl). Peak fractions were analysed by SDS-PAGE, pooled, and dialysed overnight against 4 L of buffer A. Pooled fractions from dye-ligand chromatography were filtered through a 1.2 µm syringe filter and loaded onto a 24 mL Source 15Q column equilibrated in buffer A. The column was washed with buffer A until the A280 reached a steady baseline. Rubisco was eluted with a 0 - 50% 10 column volume gradient of buffer B. Peak fractions were analysed by SDS-PAGE and the purest fractions were pooled and dialysed overnight against 4 L of buffer A. Purified Rubisco was concentrated to 10 - 20 mg/mL and filter-sterilised. Aliquots of concentrated Rubisco were flash frozen in liquid nitrogen and stored at −80 °C until use.

#### Formation of GroEL-Rubisco binary complexes

Rubisco was unfolded in unfolding buffer (50 mM HEPES-KOH, pH 7.5, 8 M urea) at 21 °C for at least 30 minutes. Binary complexes of GroEL bound to non-native Rubisco were prepared by diluting non-native Rubisco into chloride-free GroEL-containing HKM buffer (50 mM HEPES-KOH, pH 7.5, 10 mM KOAc, 10 mM Mg(OAc)2, 2 mM DTT + 1 µM GroEL tetradecamer). Unfolded Rubisco was added to 1 mL of GroEL-containing HKM buffer in five 2 µL additions. Gentle mixing and centrifugation were performed after each addition. After the fifth addition, the final concentration of Rubisco was 4 µM, a 4-fold molar excess over GroEL. The sample was incubated at 21 °C for 10 minutes with periodic mixing via gentle pipetting. Complexes were centrifuged at 16,200 RCF at 21 °C for 10 minutes to pellet aggregated protein. The presence of binary complexes was confirmed by BN-PAGE and native mass spectrometry. Binary complexes were freshly prepared for all cryoEM experiments.

#### Mass spectrometry

Samples for native mass spectrometry were exchanged into 50 mM ammonium acetate (pH 6.8) using 10 kDa cut-off Amicon Ultra centrifugal filtration units (Merck Millipore). Samples were introduced to a first-generation Waters Synapt QToF (Waters Corporation, UK) in nano electrospray gold-coated borosilicate glass capillaries (prepared in-house). Mass calibration was performed using a solution of 30 mg/mL caesium iodide (Fluka). Typical machine parameters used were: capillary 1.4 kV, sampling cone 150 V, extraction cone 4.5 V, backing pressure 7.5 mbar, trap CE 40 eV, transfer CE 10 eV, bias 88 V, source wave velocity 300 ms-1, source wave height 0.2 V, trap wave velocity 300 ms-1, trap wave height 0.2 V. Spectra were analysed using MassLynx v4.1 (Waters Corporation, UK) and Amphitrite (45). Spectra were loaded into Amphitrite using a grain size of 3 and a smoothing value of 2.

### CryoEM sample preparation and data collection

#### GroEL-Rubisco

GroEL-Rubisco was prepared and concentrated to 3.4 μM. Grids were prepared using a Chameleon instrument (SPT Labtech). We collected data from two grids frozen at different dispense-to-freeze times. Grid 1 was frozen at 1039 ms and grid 2 was frozen at 101 ms. Movies (48 frames) were collected using the EPU software on a Titan Krios transmission electron microscope (Thermo Fisher Scientific) operating at 300 keV, equipped with a Gatan K2 Summit direct electron detector in counting mode and Gatan energy filter. The defocus range was set between −1.4 and −3.0 μm, and the total exposure was 40.2 electrons/Å^2^. Images were recorded at a pixel size of 1.34 Å/pixel.

#### GroEL-ATP-Rubisco

UltrAuFoil R2/2 grids were glow discharged at 30 mA for 60 seconds using a Pelco easiGlow (Ted Pella, Inc., USA) system. ATP (3 mM) was added to GroEL-Rubisco (1 μM). Three microlitres of the mixture was applied to grids, blotted, and plunged into liquid ethane cooled by liquid nitrogen using a Vitrobot mark IV (Thermo Fisher Scientific, USA) operating at 100% humidity and 4 °C. Blot time was set to 5 s, blot force set to −10. The time between adding ATP to GroEL-Rubisco and plunge-freezing was approximately 10 sec. Movies (50 frames) were collected using the EPU software on a Titan Krios transmission electron microscope (Thermo Fisher Scientific) operating at 300 keV, equipped with a Gatan K3 direct electron detector operating in super-resolution mode and a Gatan energy filter. The defocus range was set between −1.5 and −2.7 μm, and the total exposure was 50 electrons/Å^2^. Images were recorded at a pixel size of 1.06 Å/pixel. A stage tilt of 35° was set at the start of image acquisition.

#### GroEL-ADP·BeF_3_-Rubisco

GroEL-Rubisco complexes were prepared and concentrated to 7 μM. We then added 3 mM ADP, 20 mM KF and 2 mM BeSO_4_ and incubated the sample for ten minutes. Grids were prepared using a Chameleon instrument (SPT Labtech) with a dispense-to-freeze time of 54 ms. Movies (50 frames) were collected using the EPU software on a Titan Krios transmission electron microscope (Thermo Fisher Scientific) operating at 300 keV, equipped with a Gatan K3 direct electron detector operated in super-resolution mode and a Gatan energy filter. The defocus range was set between −1.5 and −2.7 μm, and the total exposure was 50 electrons/Å^2^. Images were recorded at a pixel size of 1.068 Å/pixel.

#### GroEL-ADP·AlF_3_-Rubisco-GroES-ADP·AlF_3_

GroEL-Rubisco complexes were prepared and concentrated to 7 μM. We then added 7 μM GroES, 3 mM ADP, 20 mM KF and 2 mM KAl(SO_4_)_2_ and incubated the sample for ten minutes. Grids were prepared using a Chameleon instrument (SPT Labtech) with a dispense-to-freeze time of 54 ms. Movies (50 frames) were collected using the EPU software on a Titan Krios transmission electron microscope (Thermo Fisher Scientific) operating at 300 keV, equipped with a Gatan K3 direct electron detector operated in super-resolution mode and a Gatan energy filter. The defocus range was set between −1.5 and −2.7 μm, and the total exposure was 72 electrons/Å^2^. Images were recorded at a pixel size of 0.828 Å/pixel.

#### CryoEM image processing

The same general approach for image processing was used for all data sets. Micrograph movies were corrected for beam induced motion using Motioncorr2 (46). For movies collected in super-resolution mode using a Gatan K3 camera, micrographs were down sampled by a factor of 2 during motion correction. The CTF parameters of motion-corrected micrographs were estimated using Gctf (47). Particles were picked using the neural network particle picker included in EMAN v.2.2 (48). Particle coordinates (.box files) were imported into RELION v.3.1 (49). Particles were typically extracted from micrographs with 2 - 3 times down-sampling, giving pixel sizes of 2 - 4 Å/pixel. We used down-sampled particles for initial 2D classification, then re-extracted particles at smaller pixel sizes for 3D classification and final 3D refinements. Down-sampled particles were imported into cryoSPARC (50) and subjected to three rounds of reference-free 2D classification. Particles from featureless, noisy, or poorly resolved classes were discarded. Good particles from 2D classification were imported back into Relion using the csparc2star.py Python script (51). Subsequent image processing steps were performed in Relion v.3.1 or cryoSPARC v.3.3.1. No symmetry was applied during any step of image processing. For 3D refinements, an initial model of GroEL or GroEL-GroES was generated from a previously published cryoEM reconstruction (EMDB: 3415 and EMDB: 2325) and low-pass filtered to 30 - 60 Å.

#### GroEL-Rubisco

The image processing workflow for GroEL-Rubisco is summarised in Fig. S4. For GroEL-Rubisco, two data sets from different Chameleon grids were collected. We processed each data set separately and combined particles following 3D classification. For grid 1 (dispense-to-freeze time = 1039 ms), 175,121 particles were extracted from motion-corrected micrographs at a pixel size of 2.68 Å/pixel (384×384 pixel box, rescaled to 192×192 pixels) using Relion. Extracted particles were subjected to several rounds of iterative 2D classification in cryoSPARC. Following 2D classification, 137,331 particles were imported back into Relion, re-extracted at 1.34 Å/pixel (256×256 pixel box), and refined to 5.5 Å. The particles were then subjected to CTF refinement and refined to 4.6 Å. Particles were subjected to 3D classification using 4 classes and a 200 Å circular mask. One class (51,241 particles) displayed strong density for non-native Rubisco in the GroEL cavity and was selected. For grid 2 (dispense-to-freeze time = 101 ms), 233,763 particles were extracted from motion-corrected micrographs at a pixel size of 2.68 Å/pixel (384×384 pixel box, rescaled to 192×192 pixels) using Relion. Extracted particles were subjected to iterative 2D classification in cryoSPARC. Following 2D classification, 67,702 particles were imported back into Relion, re-extracted at 1.34 Å/pixel (256×256 pixel box), and refined to 7.3 Å. Particles were then subjected to CTF refinement, and refined to 7.0 Å. Particles were subjected to 3D classification using 4 classes and a 200 Å circular mask. One class (14,212 particles) displayed strong density for non-native Rubisco in the GroEL cavity and was selected. The selected 3D classes from the grid 1 and grid 2 data sets were combined (65,453 particles), re-extracted at 1.34 Å/pixel in a larger box (384×384 pixels), and refined to 4.4 Å. Local resolution estimation and local filtering was performed using Relion.

#### GroEL-ATP-Rubisco

The image processing workflow for GroEL-ATP-Rubisco is summarised in Fig. S5. For GroEL-ATP-Rubisco, a total of 2,267,111 particles were extracted from motion-corrected micrographs at a pixel size of 2.12 Å/pixel (256×256 pixel box, rescaled to 128×128 pixels) using Relion. Extracted particles were subjected to iterative 2D classification using cryoSPARC. Following 2D classification, 961,840 particles were imported back into Relion and refined to a resolution of 4.2 Å. In the consensus refinement, the apical domains of GroEL rings were poorly resolved. We attributed this to partial denaturation of GroEL-ATP at the air-water interface. Particles were subjected to a first round of 3D classification using eight classes. Six classes showed a poorly resolved GroEL complex resembling the consensus reconstruction. We identified two classes in which both GroEL rings were comparatively well resolved. Each of these classes displayed inter-ring asymmetry, with both a narrow ring and a wide ring, reminiscent of our previous work on GroEL-ATP (8). In class IV (105,302 particles) the wide ring was best resolved, and in class V (97,460 particles) the narrow ring was best resolved. Particles from class IV were refined to 5.3 Å, however the narrow ring was still poorly resolved. We aimed to identify subsets of particles in class IV in which the narrow ring of GroEL was intact. To do this we subtracted the better resolved wide ring from class IV particles, then performed a second round of 3D classification (skipping alignments) using eight classes. In the second round of 3D classification, classes III, IV, and VIII showed stronger narrow ring density. Particles from these 3D classes were reverted to their original non-subtracted images and refined. At a resolution of 6.5 Å, class III (13,015 particles) was the best resolved of these (others not shown) and displayed the first hints of intra-ring asymmetry. Class III particles were re-extracted at 1.7 Å/pixel (256×256 pixels rescaled to 160×160), imported into CryoSPARC and refined to 4.3 Å. The local resolution was estimated using tools in CryoSPARC and a locally filtered map was calculated. We experimented with combining different classes of particles, and with using the same particle subtraction strategy on class V from the first round of 3D classification, but these efforts led to decreased resolution.

#### GroEL-ADP·BeF_3_-Rubisco

The image processing workflow for GroEL-ADP·BeF_3_-Rubisco is summarised in Fig. S6. For GroEL-ADP·BeF_3_-Rubisco, two data sets were collected from different Chameleon grids, both with a dispense-to-freeze time of 54 ms. We processed each data set separately and combined particles following the second round of 3D classification. For grid 1, a total of 3,616,365 particles were extracted from motion-corrected micrographs at a pixel size of 3.2 Å/pixel (384×384 pixel box, rescaled to 128×128 pixels) using Relion. Extracted particles were subjected to iterative 2D classification in cryoSPARC. Following 2D classification, 2,588,967 particles were imported back into Relion and refined to 6.4 Å. Particles were subjected to 3D classification using 4 classes and a 220 Å circular mask. One class (677,383 particles) showed a well resolved GroEL complex. Particles from this class were re-extracted at 2.0 Å/pixel (468×468 pixel box, rescaled to 256×256 pixels) using Relion and refined to 4.5 Å, then subjected to CTF refinement and refined to 4.0 Å. A second round of 3D classification using a soft solvent mask was used to identify the best resolved particles in data set 1. Two classes (350,876 particles) showed a well-resolved GroEL complex and were selected for further processing. For grid 2, a total of 4,096,257 particles were extracted from motion-corrected micrographs at a pixel size of 3.2 Å/pixel (384×384 pixel box, rescaled to 128×128 pixels) using Relion. Extracted particles were subjected to iterative 2D classification in cryoSPARC. Following 2D classification, 1,792,697 particles were imported back into Relion and refined to 6.5 Å. Particles were subjected to 3D classification using 4 classes and a 220 Å circular mask. One class (592,763 particles) showed a well resolved GroEL complex. Particles from this class were re-extracted at 2.0 Å/pixel (468×468 pixel box, rescaled to 256×256 pixels) using Relion and refined to 4.4 Å, then subjected to CTF refinement and refined to 4.0 Å. A second round of 3D classification using a soft solvent mask was used to identify the best resolved particles in data set 2. Two classes (327,721 particles) showed a well-resolved GroEL complex and were selected for further processing. The selected 3D classes from the grid 1 and grid 2 data sets were combined (678,597 particles). This set of particles was re-extracted at 1.2 Å/pixel in a larger box (512×512 pixels, rescaled to 448×448 pixels) and subjected to CTF refinement and particle polishing, then imported into CryoSPARC. The set of 679k polished particles was refined to 3.3 Å, subjected to CTF refinement, then refined to 2.8 Å using local refinement in CryoSPARC. The 2.8 Å map was used to model the GroEL nucleotide binding sites during model building. However, the 2.8 Å map displayed poorly resolved density for GroEL apical domains and no density for non-native Rubisco. To identify particles with bound non-native Rubisco, we performed masked 3D classification (skipping alignments) using 4 classes and a mask that encompassed the GroEL asymmetric ring cavity. Importantly, we also increased the amplitude contrast metadata value (in the Relion .star files) of particles from 10% to 30% (mentioned briefly in ref (52)). This adjustment enabled better classification of low-resolution features such as the non-native substrate. When the amplitude contrast was set to its default value of 10%, 3D classification converged with > 99% of particles in a single class. We performed this round of 3D classification on two subsets of the data for computational efficiency. Three classes from each 3D classification job showed strong density in the asymmetric ring cavity of GroEL. We combined these classes (352,247 particles) and refined the particles to 4.0 Å. The resulting map displayed density for non-native Rubisco, but the density for the apical domains of GroEL subunits 3 and 4 was poorly resolved. We created a soft mask around the apical domains of GroEL subunits 3 and 4 and used another round of 3D classification without alignments to identify the best resolved particles. Class 1 (99,764 particles) and class 4 (102,818 particles) showed improved density for subunits 3 and 4. We refined each class separately. The two resulting reconstructions were essentially identical (reconstructions not shown). We therefore merged the particles from each class (202,582 particles) and re-extracted the combined set at 1.2 Å/pixel in a larger box (512×512 pixels, rescaled to 448×448 pixels). The re-extracted particles were then subjected to CTF refinement and particle polishing, and refined to 3.3 Å. The map displayed improved density for subunits 3 and 4. Local resolution estimation and local filtering were performed using Relion. The final map was further processed using DeepEMhancer (28). The map generated by DeepEMhancer was used to better resolve the extended apical domains of GroEL subunits 2, 5, and 7.

#### GroEL-ADP·AlF_3_-Rubisco-GroES

The image processing workflow for GroEL-ADP·AlF_3_-Rubisco-GroES is summarised in Fig. S7. For GroEL-ADP·AlF_3_-Rubisco-GroES, a total of 5,571,101 particles were extracted from 11,733 motion-corrected micrographs at a pixel size of 3.4 Å/pixel (464×464 pixel box, rescaled to 112×112 pixels) in Relion. Extracted particles were subjected to iterative 2D classification in cryoSPARC. Following 2D classification, 3,637,345 particles were imported back into Relion and refined to 6.8 Å. We split the data set into two subsets and performed 3D classification (without alignments) with a 250 Å circular mask on each subset using eight classes. GroES density varied among the 3D classes. One class from each subset showed no GroES density and resembled our reconstruction of GroEL-ADP·BeF_3_. We selected classes that displayed density for GroES (even if weak) and combined them (3,083,520 particles). We then performed a single round of 3D classification into 3 classes. One poorly resolved class contained 73.6% of the particles. Analysis by 2D classification suggested that particles belonging to the poorly resolved class were almost exclusively end views and overlapping particles (not shown). We selected the class with well-resolved GroEL-GroES features (814,317 particles), re-extracted the particles at 2.0 Å/pixel (512×512 pixel box, rescaled to 224×224) and refined them to 4.0 Å. We then performed CTF refinement and particle polishing. To identify particles with encapsulated Rubisco, we used a similar approach to our GroEL-ADP·BeF_3_-Rubisco processing workflow. We created a mask encompassing the *cis* cavity of GroEL-GroES and performed 3D classification (skipping alignments, 30% amplitude contrast) using 6 classes on two subsets of the data. One 3D class from each subset displayed a blob-like density inside the GroEL-GroES cavity. We merged these classes (30,965 particles), re-extracted particles at 0.828 Å/pixel (448×448 pixels) and refined them to 3.7 Å. The subset of 30,965 GroEL-ADP·AlF_3_-Rubisco-GroES particles was imported into cryoSPARC v.3.3.1 and subjected to a single round of 3D classification (4 classes, target resolution = 8 Å, number of O-EM epochs = 10, number of final full iterations = 5, batch size per class = 3000, initial structure lowpass resolution = 12 Å, initialisation mode = PCA, class similarity = 0) using a soft edge mask encompassing the Rubisco density. Each resulting class of particles was refined to a resolution of 4.1 - 4.2 Å. Local resolution estimation and local filtering were performed using tools within cryoSPARC.

#### Model building

For GroEL-Rubisco, we used UCSF Chimera (53) to rigid body fit the crystal structure of GroEL (PDB: 1SS8) into the locally filtered density map generated by RELION. The model was refined into the density using Isolde v.1.3 (54), followed by real space refinement using Phenix (55). Secondary structure, reference model and geometry restraints were applied during model refinement.

For GroEL-ADP·BeF_3_-Rubisco, we used the crystal structure of apo GroEL (PDB: 1SS8) as an initial model for the four substrate-contacting GroEL subunits in the asymmetric ring (subunits 1, 3, 4, and 6). These initial models were refined in Isolde using adaptive distance restraints. We applied distance restraints individually to the three domains of each GroEL subunit as follows: equatorial domain (residues 1-33, 409-525), intermediate domain (residues 134-190), and apical domain (residues 377-408, 191-376). This allowed each GroEL domain to move as a separate restrained body. Each GroEL subunit was then refined into the density. We used the crystal structure of GroEL-GroES (PDB: 1SVT) as an initial model for GroEL subunits 2, 5, and 7 and refined these models in Isolde. For the symmetric ring, we modelled a single GroEL subunit using the crystal structure of apo GroEL (PDB: 1SS8), then copied the refined model to the other six positions. The whole symmetric ring was then refined into the density. Inter-ring and inter-subunit contacts were then refined. For nucleotide binding sites, ADP, Mg^2+^, and K^+^ were placed into the density and refined using Isolde. BeF_3_ was placed into the density and refined using Coot. The structure was then iteratively refined using Coot and phenix.real_space_refine applying secondary structure, reference model, and geometry restraints. We used three maps to interpret, build and refine different parts of the model of GroEL-ADP·BeF_3_-Rubisco. First, compared to the final 3.4 Å map (202k particles), the 2.8 Å map (678k particles), determined prior to 3D classification of bound Rubisco, displayed better resolved density for GroEL nucleotide binding sites. We therefore used the 2.8 Å map (678k particles) to model ADP, BeF_3_, metal ions, and waters. Second, the map generated by DeepEMhancer was used to visualise the complex and aid rigid body fitting of GroEL subunits 2, 5, and 7, this map was not used for real space refinement. Third, the locally filtered map generated by RELION was used to refine the model using Isolde and Phenix.

For GroEL-ADP·AlF_3_-Rubisco-GroES, we used UCSF Chimera to rigid body fit the crystal structure of GroEL-GroES (PDB: 1SVT) into the locally filtered map generated by cryoSPARC. The model was refined into the density using Isolde v.1.3 and phenix.real_space_refine. ADP, Mg^2+^, and K^+^ were placed into the density using Isolde. AlF_3_ was placed into the density and refined using Coot. Secondary structure, reference model and geometry restraints were applied during real-space refinement in Phenix. To model Rubisco we used the locally filtered map of class II, here the Rubisco density had a local resolution of approximately 9 – 12 Å. We used ChimeraX (56) to perform a rigid body fit of one Rubisco monomer into the density, positioning the larger C-terminal domain (CTD) of Rubisco into the larger lobe of substrate density volume. The Rubisco monomer was flexibly fitted to the density using the Namdinator (57) web service with a map resolution of 10 Å. Further refinement of the complete model was carried out using Isolde (54) and real-space refinement in Phenix (55). Adaptive distance restraints were applied to the Rubisco monomer during refinement in Isolde.

## Acknowledgments

CryoEM data of GroEL-Rubisco, GroEL-ADP·BeF_3_-Rubisco, and GroEL-ADP·AlF_3_-Rubisco-GroES for this investigation were collected at the ISMB EM facility at Birkbeck College, University of London with financial support from the Wellcome Trust (202679/Z/16/Z and 206166/Z/17/Z). CryoEM data of GroEL-ATP-Rubisco were collected at Diamond Light Source, proposal EM20287. S.G. was supported by the BBSRC London Interdisciplinary Doctoral Programme (LIDo) (BB/M009513/1). We thank D. Houldershaw for IT support, W. Fenton for advice on protein purification, and A. Horwich for helpful comments.

## Database depositions

The cryo-EM maps and associated coordinates have been deposited in the EMDB and on the PDB: GroEL-Rubisco (EMDB: 15939, PDB: 8BA7), GroEL-ADP·BeF_3_-Rubisco (EMDB: 15940, PDB: 8BA8), GroEL-ATP-Rubisco (EMDB: 15941), GroEL-ADP·AlF_3_-Rubisco-GroES (EMDB: 15942, PDB: 8BA9), GroEL-ADP·AlF_3_-Rubisco-GroES class I (EMDB: 15943), GroEL-ADP·AlF_3_-Rubisco-GroES class II (EMDB: 15944, PDB: 8BAA), GroEL-ADP·AlF_3_-Rubisco-GroES class III (EMDB: 15945), and GroEL-ADP·AlF_3_-Rubisco-GroES class IV (EMDB: 15945).

## Author contributions

Author contributions: S.G., K.T., and H.R.S. designed research; S.G., M.C.D., and N.L. performed research; S.G. analysed data; and S.G. and H.R.S. wrote the paper.

---

**Supplementary Figure 1.**
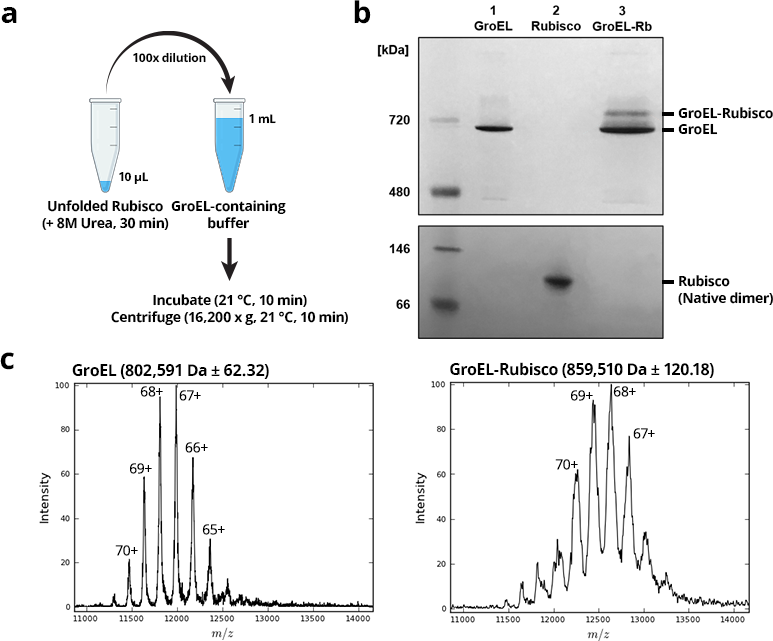
Formation of GroEL-Rubisco binary complexes. **(a)** Schematic representation of the protocol to form binary complexes. **(b)** Blue Native-PAGE of purified GroEL (lane 1), purified *R. rubrum* Rubisco in its native dimeric form (lane 2), and GroEL-Rubisco complexes (lane 3). **(c)** Native electrospray ionisation mass spectrometry (ESI-MS) of purified GroEL (left) and GroEL-Rubisco complexes (right). The charge state of each prominent peak is indicated.

**Supplementary Figure 2.**
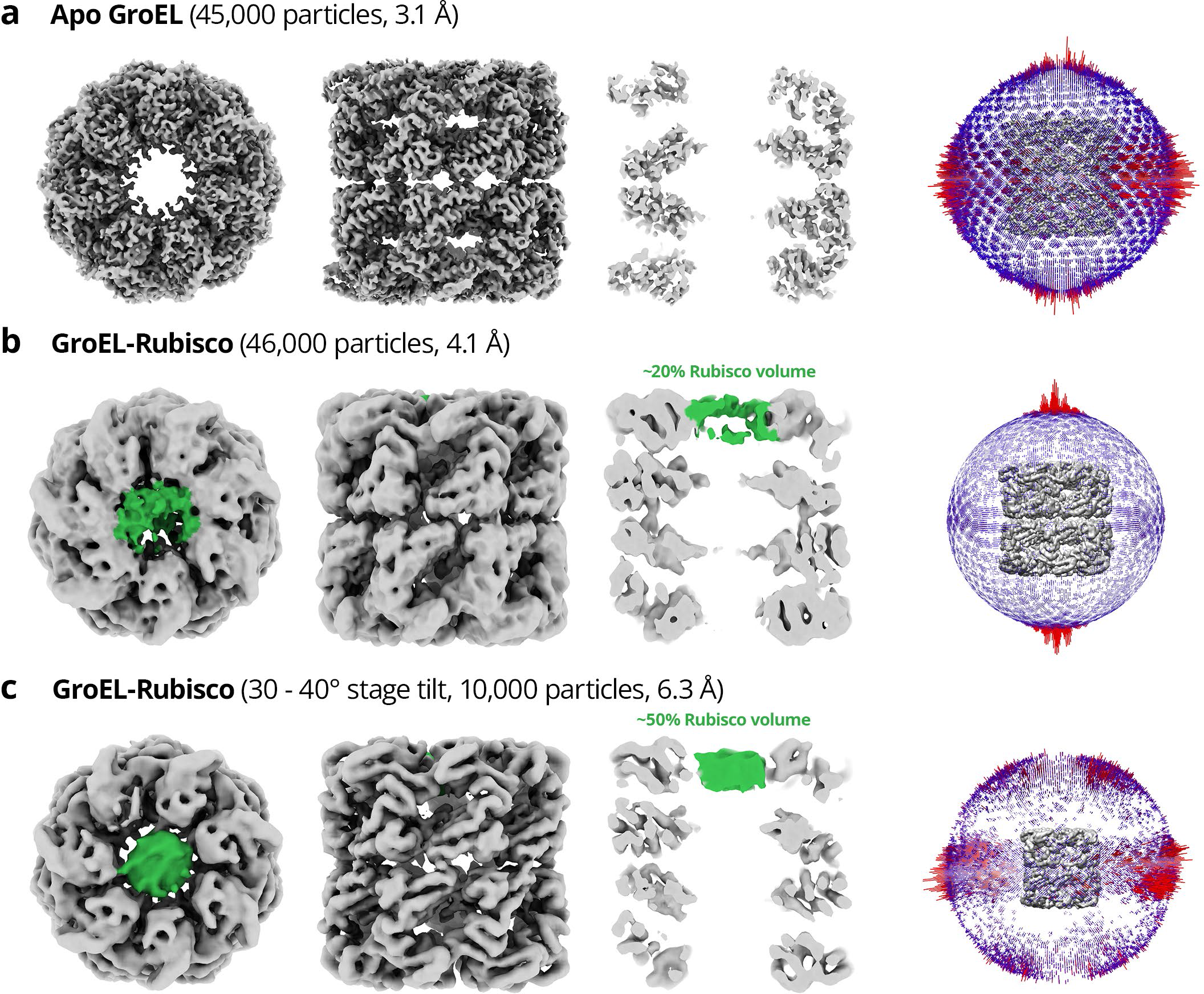
Summary of initial cryoEM experiments showing the preferred orientation of GroEL-Rubisco compared to apo GroEL. Initial cryoEM data sets were collected from grids prepared using a Vitrobot. **(a)** Final reconstruction and angular distribution of apo GroEL. **(b)** Final reconstruction and angular distribution of GroEL-Rubisco. **(c)** Final reconstruction and angular distribution of GroEL-Rubisco from cryoEM data collected employing a 30 – 40° stage tilt. Density for GroEL and Rubisco is coloured grey and green respectively. For GroEL-Rubisco complexes, the Rubisco density volume is expressed as a percentage of the theoretical volume of a folded Rubisco monomer.

**Supplementary Figure 3.**
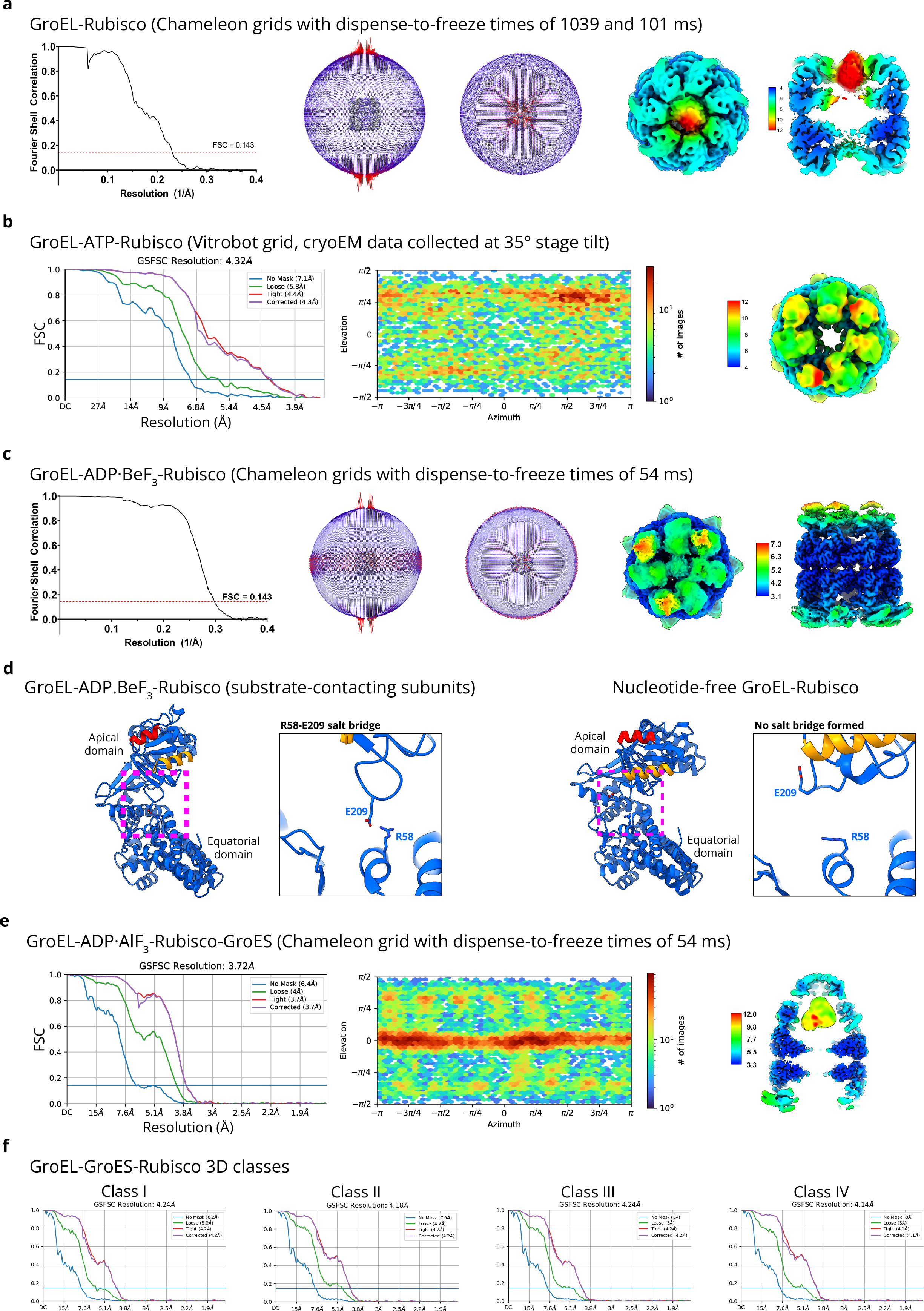
CryoEM map validation information for **(a)** GroEL-Rubisco, **(b)** GroEL-ATP-Rubisco, **(c)** GroEL-ADP·BeF_3_-Rubisco. **(d)** The R58-E209 salt bridge observed in GroEL-ADP·BeF_3_-Rubisco is not present in nucleotide-free GroEL-Rubisco. CryoEM map validation information for **(e)** GroEL-ADP·AlF_3_-Rubisco-GroES and **(f)** GroEL-ADP·AlF_3_-Rubisco-GroES 3D class refinements.

**Supplementary Figure 4.**
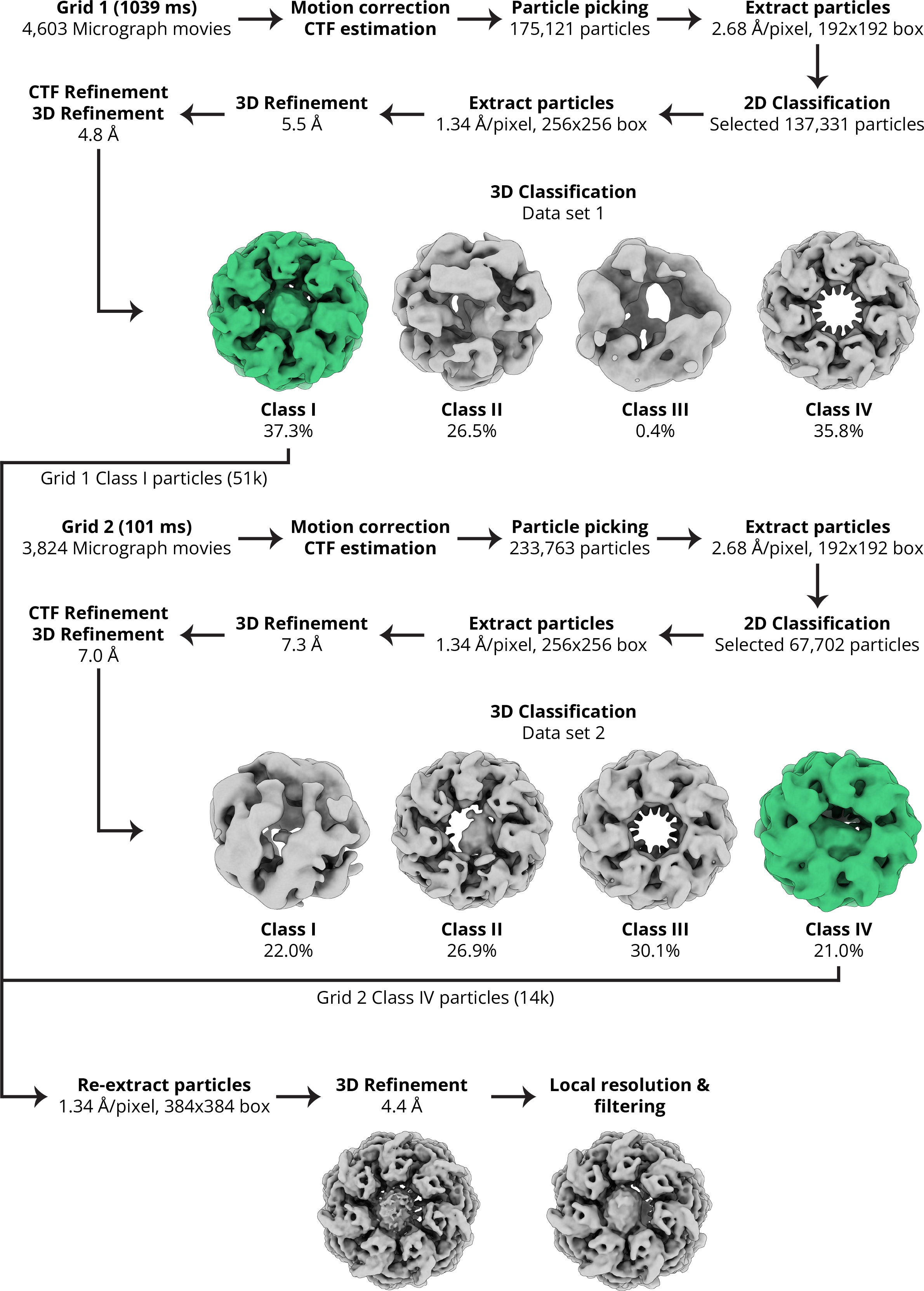
CryoEM image processing workflow for GroEL-Rubisco. Processing steps were performed using Relion v.3.1. and cryoSPARC v.3.3.1 (see methods for details). Classes selected for further processing are coloured green.

**Supplementary Figure 5.**
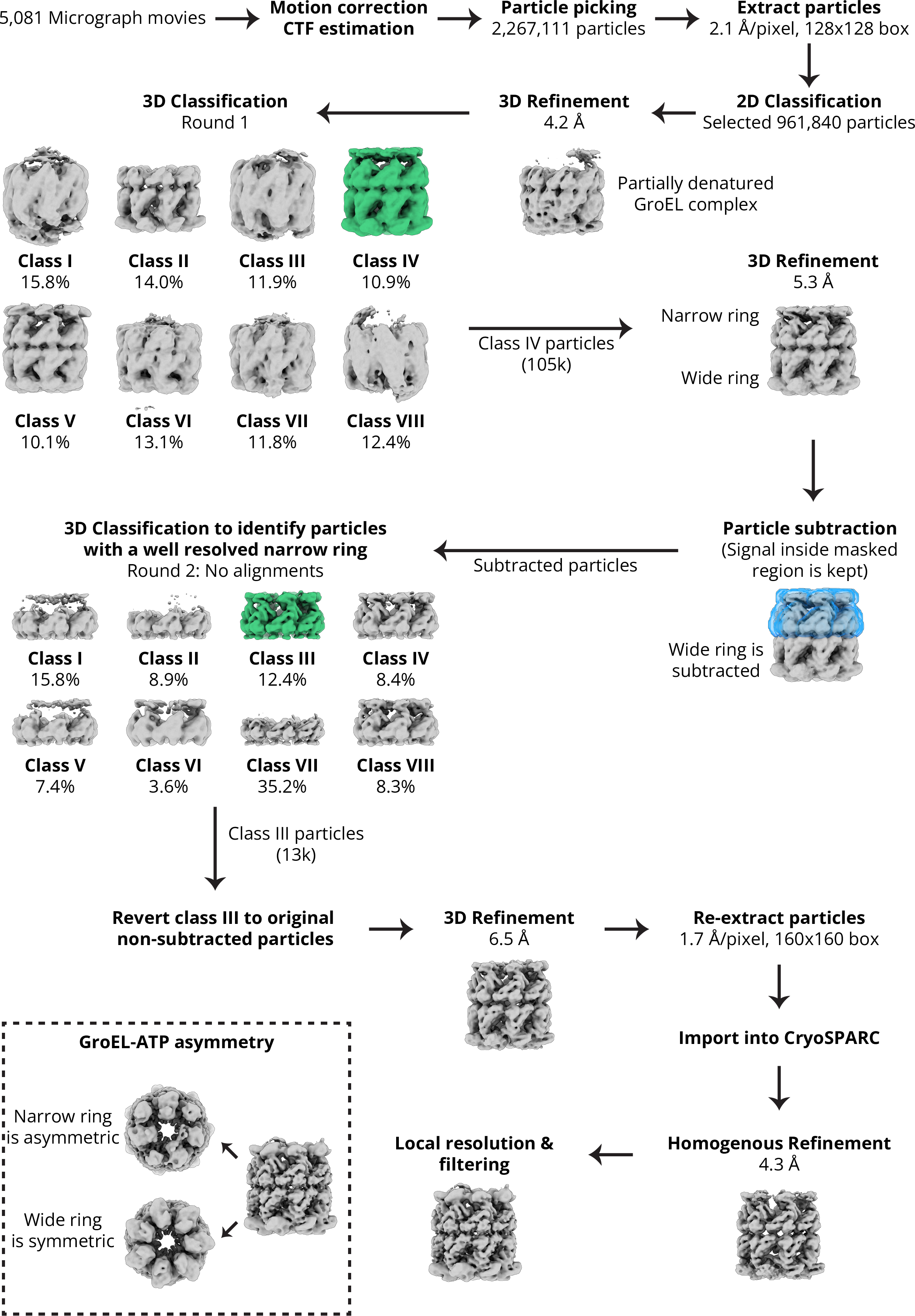
CryoEM image processing workflow for GroEL-ATP-Rubisco. Processing steps were performed using Relion v.3.1. and cryoSPARC v.3.3.1 (see methods for details). Classes selected for further processing are coloured green. Mask used during particle subtraction is coloured in blue.

**Supplementary Figure 6.**
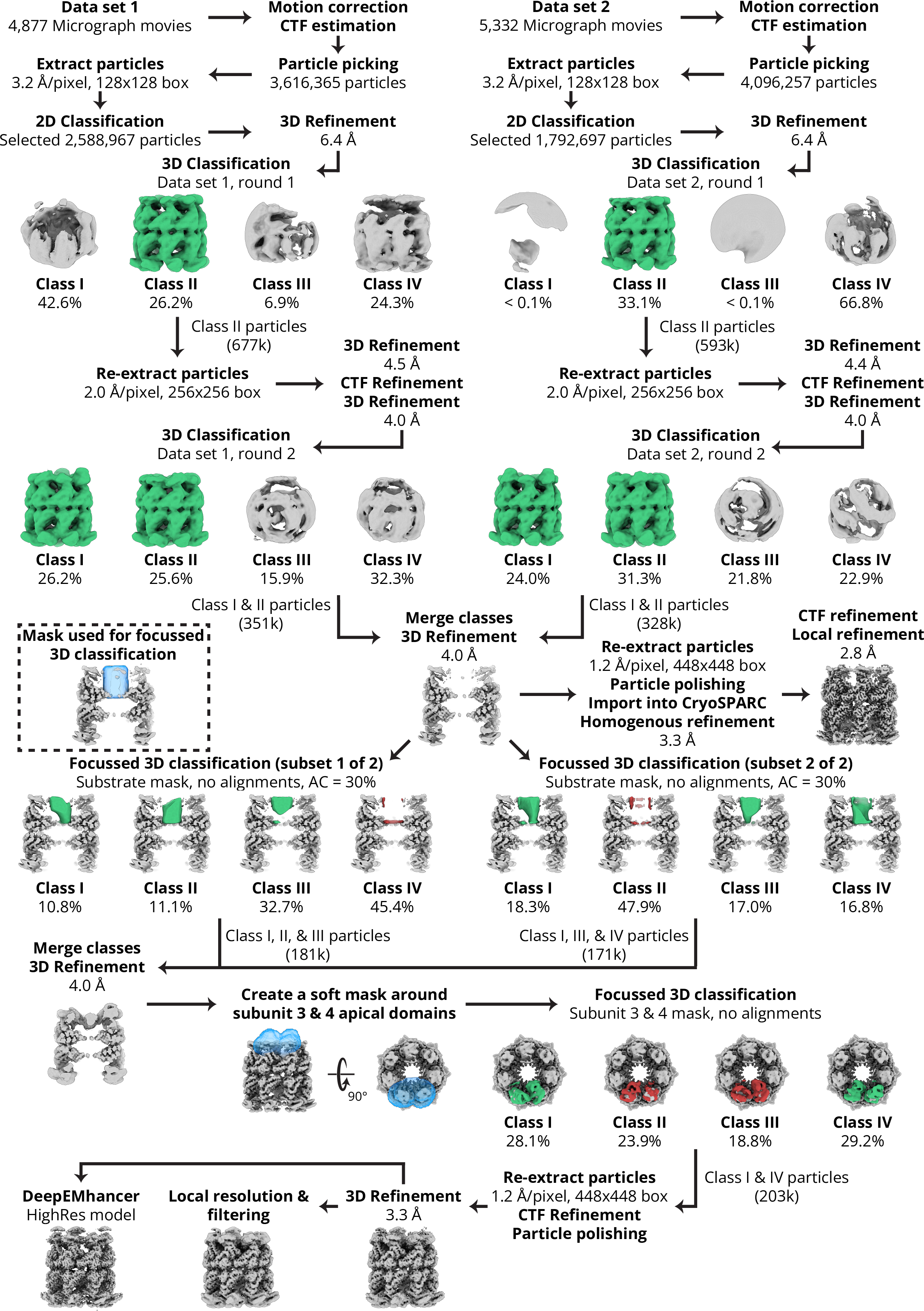
CryoEM image processing workflow for GroEL-ADP·BeF_3_-Rubisco. Processing steps were performed using Relion v.3.1. and cryoSPARC v.3.3.1 (see methods for details). Amplitude contrast abbreviated as AC. Classes selected for further processing are coloured green. In focussed 3D classification jobs, discarded classes are coloured red. Masks used during focussed 3D classification are coloured in blue.

**Supplementary Figure 7.**
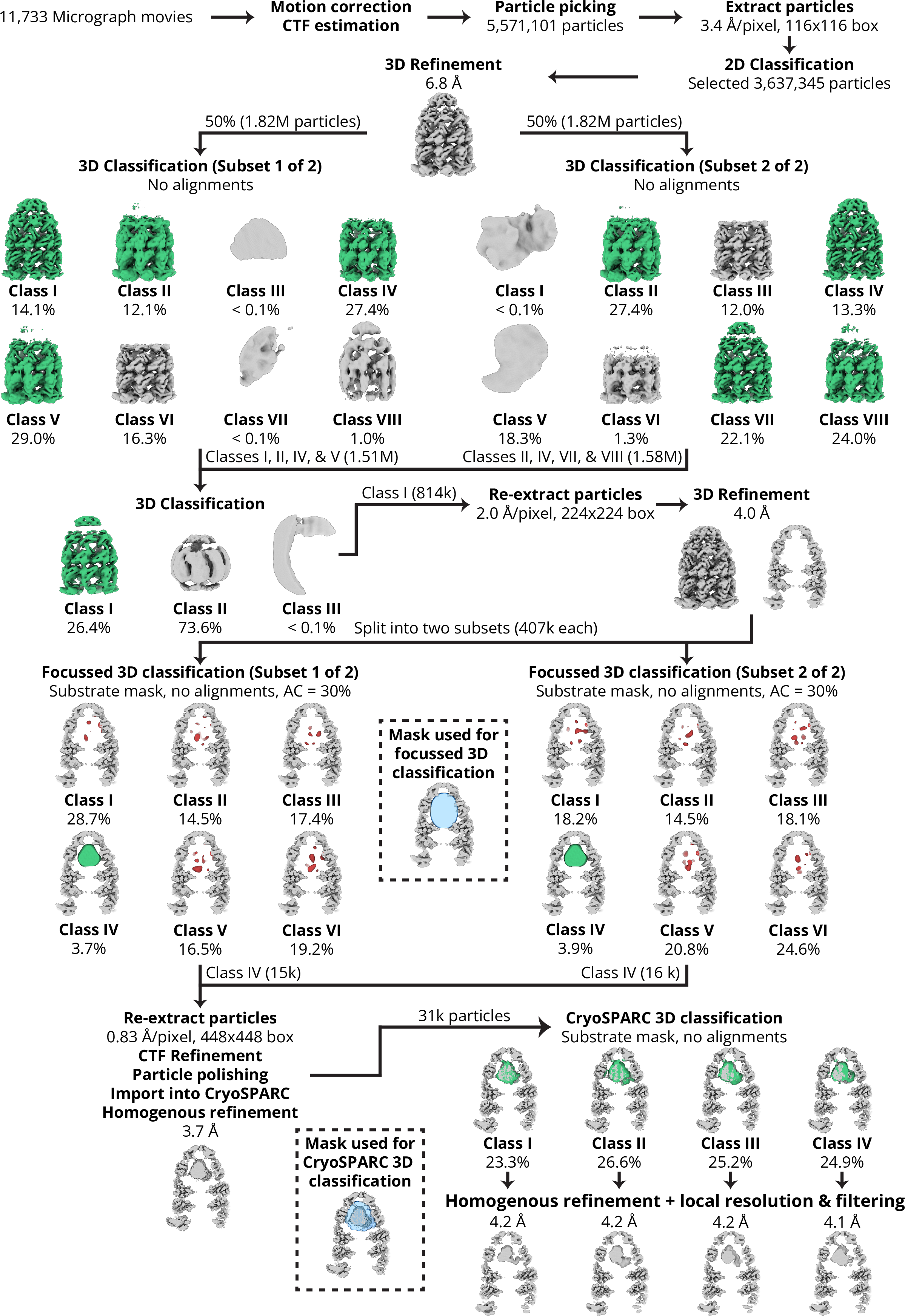
CryoEM image processing workflow for GroEL-ADP·AlF_3_-Rubisco-GroES. Processing steps were performed using Relion v.3.1. and cryoSPARC v.3.3.1 (see methods for details). Amplitude contrast abbreviated as AC. Classes selected for further processing are coloured green. In focussed 3D classification jobs, discarded classes are coloured red. Masks used during focussed 3D classification are coloured in blue.

**Supplementary Table 1.** Sample preparation details, cryoEM data collection parameters, map reconstruction and model refinement statistics.

| Sample | GroEL-Rubisco |  | GroEL-Rubisco + ATP | GroEL-Rubisco + ADP·BeF<sub>3</sub> |  | GroEL-Rubisco + ADP·AlF<sub>3</sub> + GroES |
| --- | --- | --- | --- | --- | --- | --- |
| EMDB | 15939 |  | 15941 | 15940 |  | 15944 |
| PDB | 8BA7 |  | N/A | 8BA8 |  | 8BAA |
|  | Dataset 1 | Dataset 2 |  | Dataset 1 | Dataset 2 |  |
| Sample preparation |  |  |  |  |  |  |
| Protein concentration (μM) | 3.4 | 3.4 | 1.0 | 7.0 | 7.0 | 7.0 |
| Plunge-freeze instrument | Chameleon | Chameleon | Vitrobot | Chameleon | Chameleon | Chameleon |
| Dispense-to-plunge time (ms) | 1039 | 101 | N/A | 54 | 54 | 54 |
| Data collection |  |  |  |  |  |  |
| Microscope | Titan Krios | Titan Krios | Titan Krios | Titan Krios | Titan Krios | Titan Krios |
| Voltage (keV) | 300 | 300 | 300 | 300 | 300 | 300 |
| Magnification (x1000) | 64 | 64 | 130 | 130 | 130 | 165 |
| Stage tilt (°) | 0 | 0 | 35 | 0 | 0 | 0 |
| Electron exposure (e<sup>-</sup>/Å<sup>2</sup>) | 40.2 | 40.2 | 49.1 | 50.0 | 50.0 | 72.3 |
| Defocus range (μm) | 1.4 – 3.0 | 1.4 – 3.0 | 0.5 – 2.7 | 1.5 – 2.7 | 1.5 – 2.7 | 1.5 – 2.7 |
| Detector | K2 | K2 | K3 | K3 | K3 | K3 |
| Collection mode | Counting | Counting | Super resolution | Super resolution | Super resolution | Super resolution |
| Pixel size (Å) | 1.340 | 1.340 | 1.06 | 1.068 | 1.068 | 0.828 |
| # Micrograph movies | 4603 | 3824 | 5081 | 4877 | 5332 | 11733 |
| # Particles (After 2D classification) | 137,331 | 67,702 | 961,840 | 2,588,967 | 1,792,697 | 3,637,345 |
| # Average particles per micrograph | 30 | 18 | 189 | 531 | 336 | 310 |
| Reconstruction |  |  |  |  |  |  |
| # Particles (final) | 65,453 |  | 13,015 | 202,582 |  | 8,237 |
| Map resolution (Å) (FSC threshold = 0.143) | 4.4 |  | 4.3 | 3.4 |  | 4.2 |
| Map resolution range (Å) | 4.2 – 12.0 |  |  | 3.1 – 7.1 |  | 4.0 – 12.0 |
| Map sharpening (Å<sup>2</sup>) | -130 |  |  | -60 |  | -60 |
| Model refinement |  |  |  |  |  |  |
| Model resolution (Å) (FSC threshold = 0.5) | 4.3 |  |  | 3.2 |  | 4.3 |
| Number of residues |  |  |  |  |  |  |
| Protein | 7336 |  |  | 7336 |  | 8467 |
| Ligand |  |  |  | ADP: 14<br>BeF<sub>3</sub>: 7<br>MG: 14<br>K: 14 |  | ADP: 14<br>AlF<sub>3</sub>: 7<br>MG: 14<br>K: 7 |
| Water |  |  |  | 7 |  |  |
| B-factors (Å<sup>2</sup>) |  |  |  |  |  |  |
| Protein | 162.50 |  |  | 174.29 |  | 223.68 |
| Ligand |  |  |  | 85.43 |  | 127.15 |
| Water |  |  |  | 85.06 |  |  |
| R.M.S. deviations |  |  |  |  |  |  |
| Bond lengths (Å) | 0.002 |  |  | 0.005 |  | 0.005 |
| Bond angles (°) | 0.519 |  |  | 1.072 |  | 0.847 |
| Validation |  |  |  |  |  |  |
| MolProbity score | 1.83 |  |  | 1.79 |  | 1.95 |
| Clashscore | 12.03 |  |  | 9.09 |  | 13.34 |
| Rotamer outliers (%) | 0.00 |  |  | 0.00 |  | 0.00 |
| Ramachandran plot |  |  |  |  |  |  |
| Favoured (%) | 96.43 |  |  | 95.65 |  | 95.49 |
| Allowed (%) | 3.57 |  |  | 4.35 |  | 4.45 |
| Outliers (%) | 0.00 |  |  | 0.00 |  | 0.06 |

